# A signaling network motif couples protein expression levels and intracellular localization

**DOI:** 10.64898/2026.09.04.749364

**Authors:** Diego Mazo-Durán, Antonio Juárez-Leardini, Rodrigo Torrillas, Jose María Lanza Arnaiz, Esther Serrano-Saiz, Laura Formentini, Carlos Estella, Filip Lim, David G. Míguez

**Affiliations:** Dept. de Física de la Materia Condensada, Universidad Autónoma de Madrid, 28049, Madrid, Spain; Centro de Biología Molecular Severo Ochoa, CSIC, 28049, Madrid, Spain; Instituto de Física de la Materia Condensada, Universidad Autónoma de Madrid, 28049, Madrid, Spain; Dept. de Biología Molecular, Universidad Autónoma de Madrid, 28049, Madrid, Spain

## Abstract

Most of the accepted major signaling pathways incorporate a mechanism of physical separation of active versus inactive effector proteins, in the form of dynamic nucleocytoplasmic shuttling. Here, we show that when this translocation takes place as dimers, it operates as a network motif that introduces a nonlinear coupling between expression levels of a protein and its intracellular localization. This mechanism modulates the activity of the TGF-β signaling cascade by affecting the location of its main effectors, the R-Smads, as well as in all signaling cascades analyzed that incorporate a dimerization step linked to nucleocytoplasmic shuttling, operating as a singular type of nonlinear network motif that does not require a loop of regulation and takes advantage of physical intracellular compartmentalization.

## Introduction

The interior of a living cell is a very crowded environment, and to function properly requires some spatial organization of the biological processes taking place. To achieve this, the cell utilizes a highly dynamic redistribution of proteins and other components into biological compartments. This way, proteins are actively or passively directed in and out of these cellular locations, where they interact and/or perform their functions. One of the most important cellular compartments is the cytoplasmic membrane [1], an open compartment where receptor proteins are anchored, the actomyosin cortex is organized, and proteins translocate to form complexes, interact, and initiate cascades of downstream signaling. Another main compartment is the nucleus, a closed compartment formed by the lipid nuclear membrane that mainly isolates and protects the DNA. This way, effector proteins such as transcription factors enter the nucleus through multiprotein assemblies called nuclear pore complexes (NPCs) to regulate the expression of their target genes. Similarly, messenger RNA crosses the NPCs in the opposite direction to reach the ribosomes for its translation into proteins.

Apart from this purely physical separation of processes, compartmentalization can act as an additional regulatory layer for biochemical interactions [2–3]. For instance, the cytoplasmic membrane serves as a two-dimensional scaffold that strongly decreases the activation energy of biochemical reactions by reducing the dimensionality of the domain where the interaction occurs [4]. This way, protein complex formation is more efficient when proteins and ligands diffuse and screen a two-dimensional space to find each other (compared to diffusion in 3D), accelerating the reaction rates by orders of magnitude [5–6].

Another well documented use of intracellular compartmentalization is the active or passive translocation of proteins from a large volume to a smaller one (for instance, from the cytoplasm to the nucleus). This results in a local concentration increase that directly increases the likelihood of collisions and biochemical interactions. Very often, this rate of translocation depends strongly on post-translational modifications in the proteins, such as phosphorylation, by directly or indirectly affecting the rate of nuclear import across the NPCs. This dynamic nucleocytoplasmic shuttling generates a difference in concentration between the active and inactive forms of a given protein in the nucleus and cytoplasm. Most of the major signaling pathways (NF-κB, Wnt/β-Catenin, BMP, Hh, noncanonical FGF, Calcium/NFAT, Akt, JAK/STAT, Hippo, Nuclear Receptor, and MAPK) rely on dynamic nucleocytoplasmic shuttling to modulate the expression of their corresponding target genes.

Nucleocytoplasmic shuttling is also at the core of the TGF-β (transforming growth factor beta) pathway, a highly conserved signaling cascade involved in many critical cellular processes (cell differentiation, migration, cellular dynamics and structure), and linked to many diseases (cancer, cardiovascular diseases, inflammatory and immune disorders) [7–9].

This nucleocytoplasmic translocation in the context of the canonical TGF-β activation is carried out by the Smad family of proteins and transcription factors: R-Smads (receptor-activated Smads), the Co-Smad (common mediator Smad, Smad4), and inhibitory I-Smads (inhibitor Smads, Smad6 and Smad7) [10]. In brief, Activin/Nodal ligands activate TGF-β receptors, that phosphorylate the R-Smads (Smad2 and Smad3), which dimerize and bind to one Smad4 (Co-Smad) to form active heterotrimer complexes (two R-Smads and one Co-Smad). These complexes enter the nucleus at a higher rate than monomeric R-Smads, resulting in nuclear accumulation of the active form and cytoplasmic accumulation of the unphosphorylated form (a detailed review of TGF-β signaling with a focus on the translocation process can be found in reference [11]). The last step involves the dephosphorylation and disassembly of the R-Smad/Co-Smad complex and its return to the cytoplasm, where it can be phosphorylated again by the ligand-activated membrane receptors.

Many studies focusing on TGF-β signaling rely on the nuclear accumulation of R-Smads as a widely accepted readout of TGF-β pathway activation [10, 12–16]. For instance, we have used immunostaining of Smad2 and Smad3 to monitor their translocation and interaction, as well as to show that they are simultaneously cooperating in Smad3-specific targets but antagonizing in Smad2-specific targets [17].

In the present contribution, we investigate the effect of this dynamic compartmentalization in the regulation of the activation of the TGF-β signaling cascade. We show that, in all biological scenarios analyzed, there is a clear coupling between R-Smad expression levels and protein localization, with cells with higher expression levels showing also higher nuclear ratio. We then demonstrate that this correlation is due to the differential translocation of R-Smad dimers versus monomers, acting as a regulatory network motif that shapes the dynamics, sensitivity, and response of the TGF-β pathway.

Finally, we show that all other pathways tested that rely on differential dimer translocation show the same regulation, suggesting that this motif constitutes a general mechanism that modulates dynamics and strength of signaling, similar to feedback or feedforward loops.

## Materials and methods

### Cell culture

We used standard techniques in our cell cultures for adherent cells [18], growing them in culture media (Dulbecco’s Modified Eagle Medium (DMEM) + 10 % fetal bovine serum (FBS) with 2 mM glutamine and a mixture of antibiotics available in our centre’s stock). Cells were rinsed with PBS (phosphate-buffered saline) prior to detachment with Trypsin + 0.25 % EDTA. Growing cells were kept in an incubator at 37 °C and 5 % CO_2_. When specified, upstream pathway activation was performed by adding TGF-β1 (Sigma-Aldrich, #GF346 - rehydrated in 4 mM HCl and 1 mg/ml bovine serum albumin solution) at 50 ng/mL in culture media for one hour at 37°C.

#### Generation of stable genetically modified cell lines

We used standard infection with lentiviruses protocols to generate cell lines overexpressing proteins of interest, incubating the cells for up to 48 hours. These lentiviruses were produced by HEK293 cells using transfection with Lipofectamine LTX and PLUS Reagent (Invitrogen, 15338100) of three plasmids: packing plasmid (pCMVR8.74 - Addgene, #22036), envelope expressing plasmid (pMD2.G - Addgene, #12259) and the plasmid of interest: either YFP, Smad2-YFP or Smad3-YFP. These plasmids were constructed by inserting the target sequences into a pRRL-based lentiviral transfer vector, under the control of the CMV promoter. The Smad sequences were joined to YFP by a small linker, as shown in Figure 2A.

For transfection, we seeded HEK293 cells in a P100 dish until around 80% confluent. Then, we prepared two mixtures: one of 750 μL OptiMEM (Gibco, 31985070), 5 μg of interest plasmid, 5 μg of the packaging plasmid, 2 μg of the envelope-expressing plasmid, and 24 μL PLUS Reagent; and 15 minutes later, another one of 750 μL OptiMEM and 36 μL Lipofectamine LTX. Both mixtures were combined and incubated for 15 additional minutes. Afterwards, the culture medium of the cells was removed and substituted by 2 mL of OptiMEM, and the mixture prepared before was added in drops. Next, once the cells had been incubating for 4 to 6 hours, the transfection medium was removed and substituted by DMEM + 10% FBS, which was collected up to 48 hours later. The resulting supernatant contained the lentiviruses used for infection of other cell lines.

#### Cell proliferation assay

To quantify cell proliferation, different cell lines for comparison were seeded on a 6-well culture dish so that they would reach 100% confluency at 72 hours. Images of the plates were taken every 24 hours with a Leica Flexacam C1 integrated into a Leica DM IL LED microscope. Images of the cell plates were opened in Fiji and binarized using the Threshold function, by visual inspection. The threshold was adjusted so that only the cells and not the background (dish surface) were included. Then, the percentage of the image included in the binarization corresponds to the area occupied by the cells. Doing this for all the images on the time series, we quantified the confluency rate at each time point.

#### Transient transfection of cells

Cells were seeded in a microscopy-suitable Ibidi µ-Slide 8 Well chamber and grown for 24 hours in culture media without antibiotics. Then, plasmids were introduced by using Lipofectamine 2000 (Thermo Fisher, 11668027) reagent, at a ratio of 1:3 (μg DNA to μL Lipofectamine 2000), following the standard protocol published on Thermo Fisher’s website (using OptiMEM - Gibco, 31985-062). Transfection media was removed after 4 hours, and cells were fixed 18 hours later. 1098 pcDNA GFP FKHR was a gift from William Sellers (Addgene plasmid # 9022; http://n2t.net/addgene:9022; RRID:Addgene_9022) [19]. The plasmid used for constitutive expression of the ERK2-GFP fusion protein was a gift from Piero Crespo [20].

#### Immunofluorescence

Fixation was performed using 4% PFA in culture media for 15 minutes at room temperature (RT from now on). Next, samples were washed three times with PBS 1X for 10 minutes (RT). Then, cells were permeabilized using PBT 0.5% (PBS 1X + Triton X-100 0.5%) for 20 minutes at RT and blocked with PBT-BSA (PBS 1X + Triton X-100 0.1% + 3% BSA) for 40 minutes at RT. Primary antibodies are diluted 1:200 in blocking buffer and added to samples overnight at 4°C. Afterwards, they are washed three times with PBS 1X for 10 minutes (RT), and secondary antibodies diluted 1:500 in blocking buffer are added for 60 minutes (RT). Finally, samples are treated with DAPI (Merck, 28718-90-3) diluted 1:8000 in PBS 1X, washed 2 times for 10 minutes with PBS 1X (RT), and kept at 4°C in PBS 1X until analyzed. The antibodies used for these protocols are listed below:

- Primary antibodies:

- anti-Smad2 (Cell Signalling, 5339S)
- anti-Smad3 (Abcam, ab28379)
- anti-Phospho Smad2/3 (Invitrogen, PA5-110155)
- Secondary antibody: Alexa Fluor 647 (Thermo Fisher, A-21449)

#### FRAP experiments

For FRAP experiments, cells were seeded in an Ibidi μ-Slide 8-well slide so that they were around 30% confluent 24 hours later for the experiment. Then, the sample was introduced into an *in-vivo* chamber (at 37°C and 5% CO_2_) in a Leica DMI8-CS inverted confocal microscope equipped with an HC PL APO CS2 63×/1.40 OIL immersion objective (numerical aperture 1.40, refractive index 1.518).

Pre-bleaching and post-bleaching images were taken using WLL (excitation: 514 nm, detection: 519 nm – 609 nm) at 1% power and 100% for photobleaching. The images were acquired at a resolution of 256 × 256 pixels with bidirectional scanning to increase speed. Several photobleaching scans were applied on each row, depending on the intensity of the region and adjusted by the experimenter.

For each measurement, a field with an isolated cell but containing other cells in its surroundings was chosen to perform the experiment. A region of interest (ROI) was drawn using the microscope software on the region desired to photobleach. Then, the experiment started: first, 3 images of the field were captured; second, the ROI was photobleached using 100% laser power over the area (repetitions depending on cell fluorescence intensity); and lastly, images of the full field were captured again for 5 minutes at the maximum time resolution allowed by the system (around 225 ms per frame). This sequence was fully automated by the software of the microscope.

Once we had finished the whole flow, we analyzed the data in Fiji. Three ROIs were used: one covering the photobleached region, one selecting a background area distant from the first ROI, and a third one located in another cell of the field, used as a reference. The mean intensity of the three ROIs was measured in all the frames of the experiment, including pre- and post-bleaching times. This data was used to obtain the normalized and corrected intensity of the photobleached ROI, accounting for background intensity and bleaching during image acquisition.

Let I_F_(t), I_B_(t), and I_R_(t) be the mean intensity of the photobleached, the background, and the reference cell ROIs, respectively, at a time t after photobleaching. Note that experimentally we only have discrete values for t corresponding to the frames of the image, and we set t = 0 at the first image taken right after the photobleaching process. Now, let I_F,pre_, I_B,pre_ and I_C,pre_ be the average value of the mean intensity of the photobleached, the background, and the reference cell ROIs, respectively, over all the frames taken before the photobleaching. We can now define the equation of the normalized intensity for the photobleached ROI as [21].

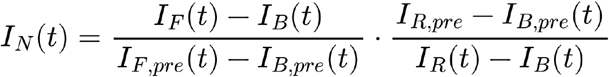

The experimental data, normalized following the equation above, was then fitted to a linear function from 0 seconds until 7 seconds using *scipy.optimize.curve_fit* (the initial regime is used to capture the dynamics without the effect of proteins returning to the photobleached region; slightly longer or shorter time windows for the fitting do not affect the measurement strongly). The intercept was set as the minimum value of I_N_(t) for each sample, while the calculated slope by the fitting was used as the photo-bleaching recovery rate.

#### Protein extraction and western blot analysis

Smad2-YFP and Smad3-YFP HaCaT proteins were extracted in lysis buffer (50 mM Tris–HCl, 150 mM NaCl, 5 mM EDTA and 1% NaP40, pH 7,5) supplemented with protease and phosphatase inhibitor cocktails. Lysates were freezed/thawed three times and clarified. Protein concentrations were determined using Bradford reagent (Bio-Rad protein assay, #5000006). The resulting supernatants were fractionated on SDS-PAGE and transferred onto polyvinylidene fluoride (PVDF) for immunoblot analysis. The primary antibodies used are anti-Smad2 (Cell Signalling, 5339S) (1:1000) and anti-Smad3 (Abcam, ab28379) (1:1000). Secondary antibody IRDye680RD goat anti-rabbit IgG (LICORBio, #926-68071) was diluted 1:10000 in 5% non-fat dried milk in Tris Buffered Saline (TBS) with 1% Tween20. An Odyssey M LICOR imaging system was used to acquire the images of the Western blot gels.

#### *In-situ* hybridization (ISH)

Fixed HaCaT Smad2-YFP and Smad3-YFP grown on top of coverslips (VWR, 631-0150) inside a 24-well plate (Falcon, 353047) were hybridized with digoxigenin-labeled riboprobes as described previously [22], with minor modifications. After fixation, cells were washed in PBS (pH 7.4), acetylated, and washed in PBS with 0.1% Triton X-100. Then, samples were incubated for 1 h at RT with hybridization buffer (50% deionized formamide (Millipore, S4117), 1X salts (pH 7.5, 10 mM Tris, 200 mM NaCl, 5 mM NaH2PO4, 5 mM Na2HPO4, 5 mM EDTA), 1X Denhardt’s (Merck, D2532), 10% dextran sulphate (Merck, 4911), tRNA 1 mg/ml (Merck, R6625)) in a humified chamber with 5X SSC (pH 4.5, 750 mM sodium citrate, 75 mM NaCl) and 50% deionized formamide (Millipore, S4117). Then, cells were hybridized overnight at 72°C with riboprobes (1:1000 in hybridization buffer).

After hybridization, samples were washed for 90 min in 0.2X SSC at 72°C, and blocked in 2% blocking solution (Roche, 11096176001) in 1X MABT (pH 7.5, 100 mM maleic acid, 150 mM NaCl, 0.1% Tween-20 (Merck, P1379)) for 1 h at RT. After blocking, they were incubated overnight at RT with anti-DIG antibody conjugated with alkaline phosphatase (1:5000, Roche, 11093274910) in MABT. After extensive washing in TBST (100 mM Tris, 150 mM NaCl, 0.1% Tween-20) at pH 7.5 and then in TBST at pH 9.5, the alkaline phosphatase activity was detected at RT by using 100 μg/ml 4-nitro blue tetrazolium chloride (Roche, 11383213001) and 175 μg/ml 5-bromo-4-chloro-3-indolyl phosphate p-toluidine salt (Roche, 11383221001) in NTMT (pH 9.5, 100 mM Tris, 100 mM NaCl, 50 mM MgCl2, 0.1% Tween-20).

Cells were mounted on microscopy slides (VWR 631-1550) with Prolong (Thermo Fisher, P36984), after performing an immunofluorescence (as described above) using anti-GFP primary antibody (Abcam, ab13970) and Alexa Fluor 647 (Thermo Fisher, A-21449) secondary antibody. This step was necessary for YFP visualization, as the in-situ protocol destroys YFP chromophore fluorescence.

Riboprobes were generated de novo in this study for the CDKN1A gene (cloned in pSC-A-Amp/Kan - Strataclone, 240205).

*In situ* hybridization combined with immunofluorescence images were acquired using a CMOS camera mounted on a fluorescence microscope. Immunofluorescence signal was captured using standard DAPI and RFP excitation/emission filters. Images were collected as 3-channel composites at a resolution of 1360 × 1024 pixels. To optimize signal quality and prevent saturation, the light source intensity and camera exposure times were adjusted manually by the experimenter on each row/region, depending on the local signal intensity.

#### Drosophila Fat Body

Third instar larvae of a strain that expresses GFP-tagged Stat92E from the native promoter (Bloomington #38670) were dissected in PBS and fixed in a solution of 4% paraformaldehyde, 0.1% deoxycholate, and 0.1% Triton X-100 in PBS for 30 min at room temperature. We used Phalloidin (TRITC) (Sigma-Aldrich, #P1951) to stain the actin cytoskeleton to label the cell membranes and DAPI (Merck) to stain nuclei. The fat body were dissected and mounted in Vectashield (Cat# H-1000 RRID: AB_2336790) for confocal analysis.

### Chick embryo neural tube immunofluorescence

Fertilized White-Leghorn eggs (Granja Gibert, Tarragona, Spain) are kept at 4°C until incubation begins, then they are incubated at 38.5°C in an incubator with airflow and an atmosphere of 70% humidity. Chick embryos were staged according to Hamburger and Hamilton [23]. Around 120 hours later, the eggs are opened, and the embryos are extracted and cleaned in PBS. Extirpation of limbs, heart, and tail takes place before fixation for 2 hours at 4°C with PBS + 4% PFA. Then, they are rinsed with PBS and embedded in a gel (distilled water with 10 %(m/v) saccharose and 5% (m/v) agarose) for slicing in a vibratome (with a 2 mm oscillation amplitude, an advance speed of 4 mm/s, and a section thickness of 70 μm).

Then, slices are placed in an M24 cell culture well with PBS, where immunofluorescence is performed. The slices are washed and permeabilized, treated twice for 5 minutes each with PBT (PBS + Triton X-100 0.1%) at RT. Blocking for 30 minutes with 1% BSA in PBT (RT) comes before primary antibody incubation for 2 hours at RT, diluted 1:500 in PBT + 1% BSA. Next, samples are washed twice for 5 minutes (RT) with PBT + 1% BSA, and the secondary antibody is then applied for 2 hours at RT, diluted 1:500 minutes in PBT + 1% BSA, ensuring the well is covered. Next, we perform two washes with PBT for 5 minutes (RT), keeping the well covered. Afterwards, samples are again washed for 5 minutes (RT) with PBS 1X and 1:5000 DAPI (Merck, 28718-90-3). Slices are finally mounted with Mowiol (Merck, 81381) on microscopy slides (VWR, 631-1550) with a coverslip (VWR, 631-0121). The antibodies used for these protocols are listed below:

- Primary antibodies:

anti-Smad2 (Cell Signaling, 5339S)
anti-Smad3 (Abcam, ab28379)
- Secondary antibody: Alexa Fluor 555 (Thermo Fisher, A-31572)

### Embryonic bodies immunofluorescence

Human cortical embryonic bodies (AG08CS (XY) at day 30) were a kind gift from Leonardo Beccari’s laboratory. We fixed them by removing the culture media and rinsing the bodies with PBS 1X, and adding PBS 1X + 4% PFA for one hour at RT. Samples are then embedded in a gel (distilled water with 10 % (m/v) saccharose and 5 % (m/v) agarose) for slicing in a vibratome (with a 2 mm oscillation amplitude, an advance speed of 1 mm/s, and a section thickness of 70 μm).

For the immunofluorescence assay, after a single 10-minute wash in PBS 1X, the sample is permeabilized and washed three times for 20 minutes with PBT 0.5% (PBS 1X + Triton X-100 0.5%). Blocking buffer (PBT 0.5% + 1% BSA + 5% FBS) is later used to block the samples for one hour, and it is the solvent for the rest of the reactants. The bodies are incubated with primary antibodies 1:500 overnight at 4°C, followed by three 15-minute washes with PBT 0.5%. Then, secondary antibodies at 1:1000 are added, and samples are incubated for two hours at RT.

Finally, after washing three more times for 15 minutes with PBT 0.5%, we incubate the samples with 1:5000 DAPI (Merck, 28718-90-3) in PBT 0.5% for 30 minutes at RT, concluding the staining with two 10-minute washes in PBT 0.5%, followed by two 10-minute washes in PBS 1X. Slides are mounted with Mowiol (Merck, 81381) on microscopy slides (VWR, 631-1550) with a coverslip (VWR, 631-0121). The antibodies used for these protocols are listed below:

- Primary antibodies:

anti-Smad2 (Cell Signaling, 5339S)
anti-Smad3 (Abcam, ab28379)
- Secondary antibody: Alexa Fluor 488 (Thermo Fisher, A-21206)

### Microscopy imaging and image analysis

Unless stated otherwise, images were acquired using a Leica DMI8-CS inverted confocal microscope equipped with an HC PL APO CS2 63×/1.40 OIL immersion objective (numerical aperture 1.40, refractive index 1.518). Fixed samples were excited using a 405 nm diode laser and a White Light Laser (WLL) system. Fluorescence emission was collected across four spectral detection channels depending on the sample: DAPI (excitation: 405 nm, detection: 415–477 nm), Alexa 488 (excitation: 499 nm, detection: 509–568 nm), Alexa 555 (excitation: 553 nm, detection: 563–620 nm), and Alexa 647 (excitation: 650 nm, detection: 660–750 nm). Pinholes were set to 1.0 Airy units for all channels to ensure optimal optical sectioning. Images were captured at 512 × 512 pixel resolution in 16-bit format, operating at a scanning speed of 600 Hz and a pixel dwell time of 0.96 μs without line or frame averaging. Large-area images were acquired by tile scanning and automatically stitched using the mosaic merging algorithm within the microscope’s control software (Leica Application Suite X, LAS X).

Image analysis and processing were performed on Fiji (ImageJ) software, on the latest version available.

#### Quantification of the cell cultures infection rate

Images of HaCaT cells infected with Smad2-YFP and Smad3-YFP after nuclear staining with DAPI were used to estimate the transfection rate. The nuclei channel was segmented using Fiji’s built-in watershed algorithm, generating a set of ROIs (one for each cell). Then, using the position of the ROIs in the YFP channel, they were classified into two groups: one with YFP signal inside (transfected cells) and another with no YFP signal inside (non-transfected cells). This allowed us to obtain the transfection ratio for each cell type.

#### Fluorescence intensity analysis

Once images were taken, their analysis was continued in Fiji. First, a Gaussian blur filter was applied in order to reduce noise. The radius varied according to the analyst’s criteria, being just big enough to smooth the area to quantify, typically between 0.5 and 2.0. This ensured that the selected areas for quantification were more homogeneous and less dependent on their exact location within the nucleus/cytoplasm. Then, the “Oval” tool was used to draw an ROI in the nucleus of the cell using the DAPI channel as a reference and avoiding the nucleolus, approximately covering 40% of it; next, the mean intensity of the ROI in the overexpressed protein channel and the immunofluorescence channel (if any) was measured by the “Measure” tool integrated in Fiji. The same procedure was followed after drawing an ROI in the cytoplasm of the cell (approximately the same size as the one used in the nucleus), and this process was repeated for several cells in the image, deliberately including a wide range of overexpressed protein levels.

After quantifying nuclear and cytoplasmic mean intensity for each cell with Fiji, we summed these two quantities to obtain the total mean intensity of the cell. Then, dividing the nuclear by the total mean intensity, we obtained the desired nuclear ratio.

### Data analysis, numerical simulations, plots and fittings

All numerical and data analysis were performed using Python 3.13, using numpy for mathematical operations. Numerical simulations of the ODE systems were made with *scipy.integrate.odeint* solver. Plots were made with matplotlib. Statistical differences on violin plots were calculated using an unequal variance t-test with the function *scipy.stats.ttest_ind* (a Welch’s t-test), and the significance code used in plots is indicate that p-value (p) is: p < 0.001 (***), 0.01 ≥ p > 0.001 (**), 0.05 ≥ p > 0.01 (*) or 0.1 ≥ p > 0.05 (n.s.).

Nuclear ratio curves predicted by our model were fitted to their experimental data using the bootstrap method as explained as follows.

Let x_exp_ and y_exp_ be the experimental data for independent and dependent variables, respectively. Each vector has N elements, so that each data point is identified with the index i = 1, 2,…, N. We want to fit this data to an analytical function f(x, p) that has one parameter p, whose value we want to obtain. For simplicity in the explanation, we will just use one parameter, but p could also be a vector with a number of different parameters. In our specific case, p will be k_1_. We define the error function as

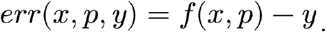

Using least squares fitting (*scipy.optimize.least_squares*), we obtain p_fit_, the value of the parameter from the fitting, minimizing err(x, p, y) for each pair (x_exp_, y_exp_). We calculate the standard deviation of the residuals of the fit as

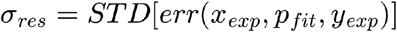

using *numpy.std*. Now, for estimating the goodness of fit, we generate M different datasets, each one labeled with the index m = 1, 2,…, M. The data points for the new datasets are:

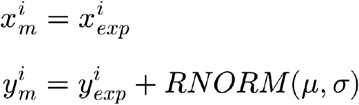

with RNORM(μ, σ) a function that returns a random number with a normal distribution with mean μ = 0 and standard deviation σ = σ_res_ (*np.random.normal*). For each dataset, we perform the least squares fit to obtain M new values of p*_fit_*. Defining μ_p_ and σ_p_ as the mean value and the standard deviation of all p_fit_, respectively, the final value with 95% CI is μ_p_ ± 1.96σ_p_. This is true because we assume that all the p_fit_ values are normally distributed, by the Central Limit Theorem.

As this is a method involving randomly generated numbers, results of σ_p_ vary slightly each time we perform the analysis. To overcome this issue, we used a big number of randomly generated datasets (M = 1000), so that results are reliable.

## Results

### Nuclear translocation of the R-Smads correlates with R-Smad expression in different biological contexts

As mentioned previously, the nuclear accumulation of the R-Smads is a well-established real-time readout of TGF-β pathway activity. Here, we apply this approach to identify and quantify specific regions with differential activation of the TGF-β signaling in different tissues. Figure 1A presents transversal sections of the developing chick spinal cord at 72 hpf (see Methods) stained with DAPI (first column) and immunofluorescence against Smad2 and Smad3 (second column) acquired with a confocal microscope. Both R-Smads show a clear expression pattern mainly restricted to the subventricular zone, occupied by the cycling progenitors. Inside this region, Smad2 (first row) exhibits a dorsoventral expression pattern with increased levels in the middle region of the spinal cord, and lower expression above (dorsal) and below (ventral). On the other hand, Smad3 (second row) exhibits a clear apicobasal pattern, with higher expression in the region corresponding to the transition zone between cycling progenitors and differentiated neurons. The third and fourth columns show a zoom at cellular resolution to compare regions with different overall R-Smad intensity levels. In both cases, regions in the tissue with lower Smad2 or Smad3 levels (blue square) show a more cytoplasmic distribution of the staining than regions where expression levels are increased (red square), an indication of higher TGF-β pathway activity. Interestingly, these regions of high TGF-β activity do not correlate with the regions of high expression of TGF-β ligands reported for the developing spinal cord (roof plate, notochord and floor plate) [24–26], suggesting that other processes may be modulating the activation of the TGF-β pathway, or the localization of its downstream effectors.

**Figure 1.**
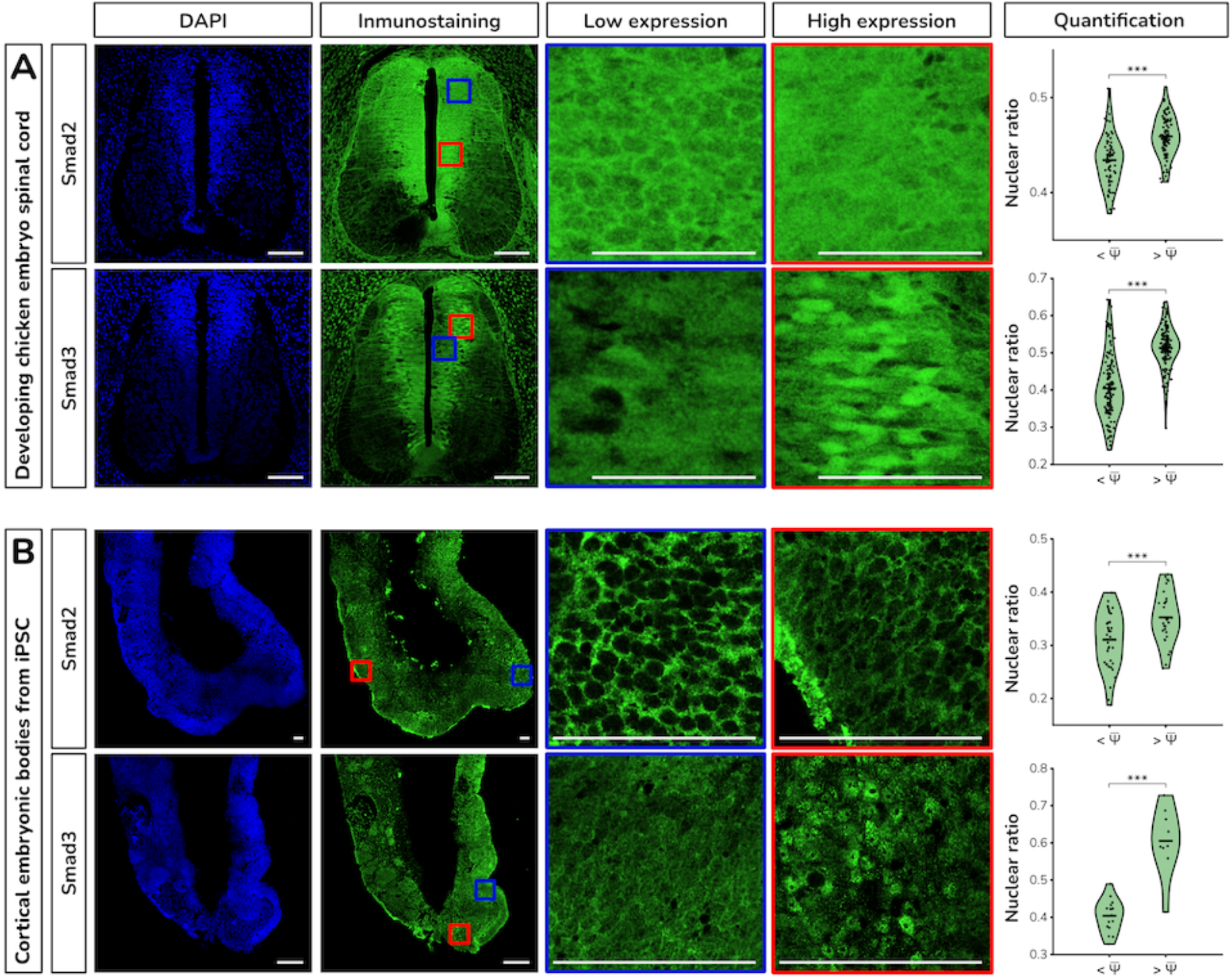
(A-C) Confocal images of biological tissues stained with DAPI (first column) and immunofluorescence against Smad2 and Smad3 (second column) for (A) transversal confocal sections of the developing chicken spinal cord at 72 hpf (scale bars: 100 µm); (B) Embryoid bodies generated from human IPS cells at 30 hours (scale bars: 100 µm). The horizontal black line marks the mean value of the nuclear ratio of each subset.

Next, we use the DAPI and immunofluorescence channels to quantify the nuclear ratio (intensity in nuclei divided by intensity in nuclei plus intensity in cytoplasms, see Methods) of both R-Smads as a measure of the TGF-β pathway at the single-cell level [17]. The right panels plot the values of the nuclear ratio at single cell resolution, clustered based on total R-Smad expression (above and below the average Smad2 or Smad3 level). In both scenarios, R-Smad levels below the average tend to have a lower nuclear ratio than cells expressing levels above the average value, with statistical significance.

Next, we investigate if this correlation between expression level and nuclear ratio is specific to the developing spinal cord, or it is a feature present in other developmental systems where TGF-β has been shown to have a central role, such as mammalian brain development. Since R-Smad antibodies fail to penetrate cells in brain tissues, we turn to cortical embryonic bodies (EBs) from human iPSC cells (see Methods), where members of the TGF-β pathway are highly expressed [27] and govern axon specification [28] among other processes. Representative images of cryostat sections acquired using a confocal microscope and stained for DAPI and immunofluorescence against Smad2 or Smad3 are shown in Figure 1B. The outer layer of the EBs reproduces the same spatial organization of the embryonic neural tube and the ventricular zone of the developing brain, as a polarized neuroepithelium. Again, visual comparison of average intracellular organization shows Smad2 as more cytoplasmic than its partner Smad3, and regions with higher expression levels (red square) have a higher nuclear accumulation than regions of lower R-Smad expression (blue square). Quantification of the nuclear ratio in individual cells clustered based on total expression levels (right panels) shows once more that cells with Smad2 or Smad3 levels below the mean level of expression have a lower nuclear ratio than cells with expression levels above the mean, with strong statistical significance.

In conclusion, despite their highly similar homology, structure, and activation mechanism, Smad2 and Smad3 present different expression patterns in all systems analyzed. In addition, Smad3 tends to have a higher nuclear accumulation than Smad2 in all three systems. Also, cells expressing higher levels of Smad2 or Smad3 always have a higher nuclear ratio than cells expressing lower levels.

### Fluorescently tagged R-Smads overexpression shows high variability, reproducing the response and dynamics of the endogenous R-Smads

To further investigate the correlation of R-Smad expression levels with their intracellular distribution, we turn to a more controlled *in vitro* approach, based on lentiviral infection of Smad2 and Smad3 fused to YFP for direct visualization. The scheme of the plasmids and the lentiviral infection process is illustrated in Figure 2A-B. Images of the HaCaT cell line (human keratinocytes, see Methods) infected with these constructs are shown in Figure 2C, showing a similar distribution to the endogenous proteins in tissues, with Smad3 being more nuclear than Smad2 on average.

**Figure 2.**
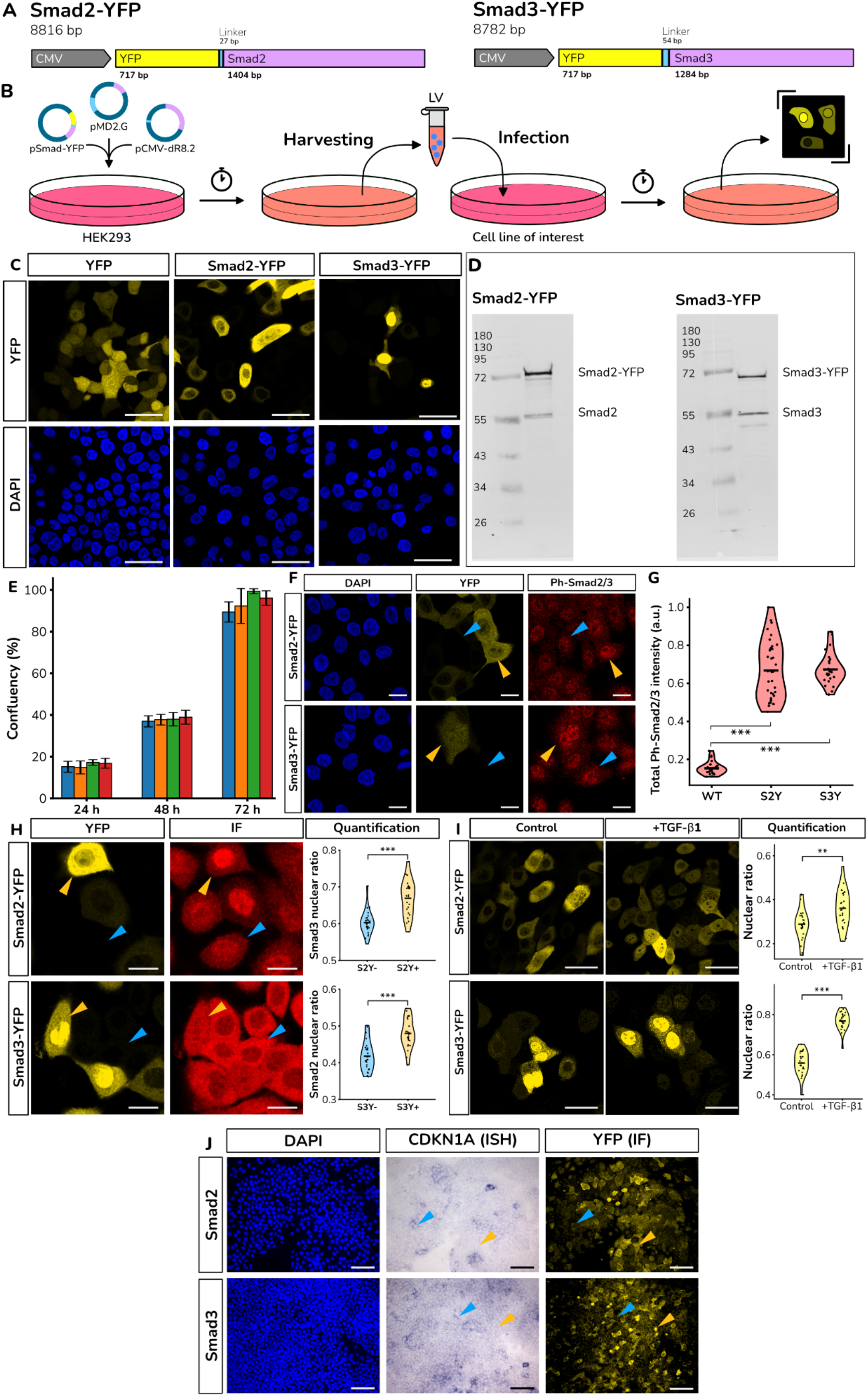
(A) Scheme of the fluorescently tagged R-Smads in lentiviral vectors. (B) Scheme of the lentiviral infection experiment. (C) Images of HaCaT cells expressing YFP, Smad2, and Smad3 (scale bars: 50 µm). (D) Comparison between endogenous R-Smad levels and exogenous levels via Western Blot against Smad2 and Smad3. (E) Quantification of growth rate of control cells, cells overexpressing YPF, Smad2-YFP, and Smad3-YFP showing similar cell cycle length. (F) Snapshots of cells showing that high Smad2-YFP and Smad3-YFP levels correlate with higher levels of phosphorylation (scale bars: 20 µm). (G) Violin plots showing higher phosphorylation levels in cells expressing Smad2-YFP and Smad3-YFP. (H) Snapshots of cells expressing Smad2-YFP and immunostained (IF) against Smad3 (upper row) and cells expressing Smad3-YFP and immunostained (IF) against Smad3 (bottom row) to show that the presence of the exogenous protein affects the localization of the endogenous one (scale bars: 20 µm). (I) Snapshots before and after TGF-β activation, showing higher nuclear localization of the exogenous proteins after activation (scale bars: 50 µm). (J) RNA expression levels of CDKN1A, a well-characterized target of Smad2 and Smad3 in HaCaT cells estimated by in situ hybridization, show no correlation with expression levels of Smad2-YFP and Smad3-YFP, suggesting that our proteins are not able to drive the expression of transcriptional targets of the pathway (scale bars: 100 µm).

Comparison of the expression level of the exogenous versus fluorescently tagged R-Smads is performed by Western blot using antibodies against Smad2 and Smad3 (Figure 2D). Results normalized by the average number of cells infected in the culture (98.87% for Smad2 and 61.65% for Smad3) suggest an average increase of 3.08x for Smad2-YFP and 1.42x for Smad3-YFP.

Overexpression of fluorescently tagged proteins has been shown to potentially result in higher oxidative stress and interfere with the normal response of the cells [29–30]. To estimate the effect of the overexpression, we quantified the growth rate of infected and uninfected cells over 4 days [31–32]. Results shown in Figure 2E show that cells expressing Smad2-YFP or Smad3-YFP grow normally at the same rate as control cells and cells overexpressing YFP, suggesting that R-Smad or YFP overexpression at these levels does not interfere with the normal growth of the cells *in vitro*.

Next, to test if the fusion proteins are being phosphorylated in response to upstream pathway activation, we performed immunofluorescence against the phosphorylated form of Smad2 and Smad3 in Smad2-YFP or Smad3-YFP expressing cells. Figure 2F shows higher phosphorylation in cells with higher fluorescence, confirmed by the quantification in Figure 2G.

Next, to test if the exogenous proteins behave as expected in response to pathway activation, we cultured cells in the presence of TGF-β1 ligand. Figure 2I shows a higher nuclear localization of the exogenous proteins after activation, suggesting that the fluorescently tagged proteins behave similarly to the endogenous R-Smads.

After validating that the fluorescently tagged R-Smads get phosphorylated, dimerize, and translocate to the nucleus, we investigated their effect on the expression of downstream targets characteristic of TGF-β signaling. To do this, we purified RNA from HaCaT cells and designed and generated probes for CDKN1A, a well-characterized target of both Smad2 and Smad3 [33]. This is then used to perform an *in situ* hybridization (ISH, see Methods) followed by immunofluorescence against YFP (to detect cells overexpressing our exogenous proteins. Figure 2J illustrates the staining over a region of cells, showing no correlation between cells expressing higher levels of CDKN1A mRNA and cells expressing higher levels of Smad2-YFP or Smad3-YFP. This lack of correlation suggests that the fluorescently tagged R-Smads have minimal transcriptional activity in driving the expression of CDKN1A. One possibility is that the presence of the YFP in the N-terminus prevents them from recruiting the required cofactors for transcriptional activity. In addition, this lack of transcriptional activity is consistent with their minimal or null effect in cell growth and cell proliferation, and suggests that they may be used as a highly innocuous dynamic readout of TGF-β pathway activation.

In conclusion, detailed characterization of the exogenous R-Smads illustrates that they recapitulate a similar intracellular organization as the endogenous proteins; they are phosphorylated, dimerize, and enter the nucleus following upstream TGF-β1 ligand stimulation similarly to the endogenous proteins. On the other hand, despite their high expression levels compared to the endogenous proteins, they have a minimal effect on the growth rate and cell cycle of the host cells, and they are not transcriptionally active. These characteristics make them ideal as a ideal tool for a real-time *in vivo* readout of the activity of TGF-β signaling

### The nuclear ratio of exogenous R-Smads *in vitro* correlates with R-Smad expression levels

In the previous section, we have shown that the lentiviral infection of exogenous R-Smads provides a reliable and direct readout of the dynamics of TGF-β signaling in a highly controllable *in vitro* environment where all cells receive the same upstream TGF-β stimulation. Here, this tool is used to study the changes in intracellular localization of the R-Smads induced by changes in their expression level. Snapshots of HaCaT cells cultured *in vitro* two days after infection with YFP, Smad2-YFP, or Smad3-YFP are shown in Figure 3A-C. To visually illustrate the differences in nuclear ratio, the same image is reproduced twice with contrast settings adjusted to visualize low-expressing cells (left panel) and highly expressing cells (right panel). Visual comparison shows an increase in nuclear ratio for highly expressing Smad2-YFP and Smad3-YFP, but not for YFP (blue arrows point to individual cells expressing low fluorescence, and orange arrows point to individual cells expressing high fluorescence). Quantification of nuclear ratio for cells expressing levels below and above the mean (right panel) shows again a statistically significant difference in nuclear ratio between cells expressing below and above the average for both R-Smads, but not for the cells expressing YFP only.

**Figure 3.**
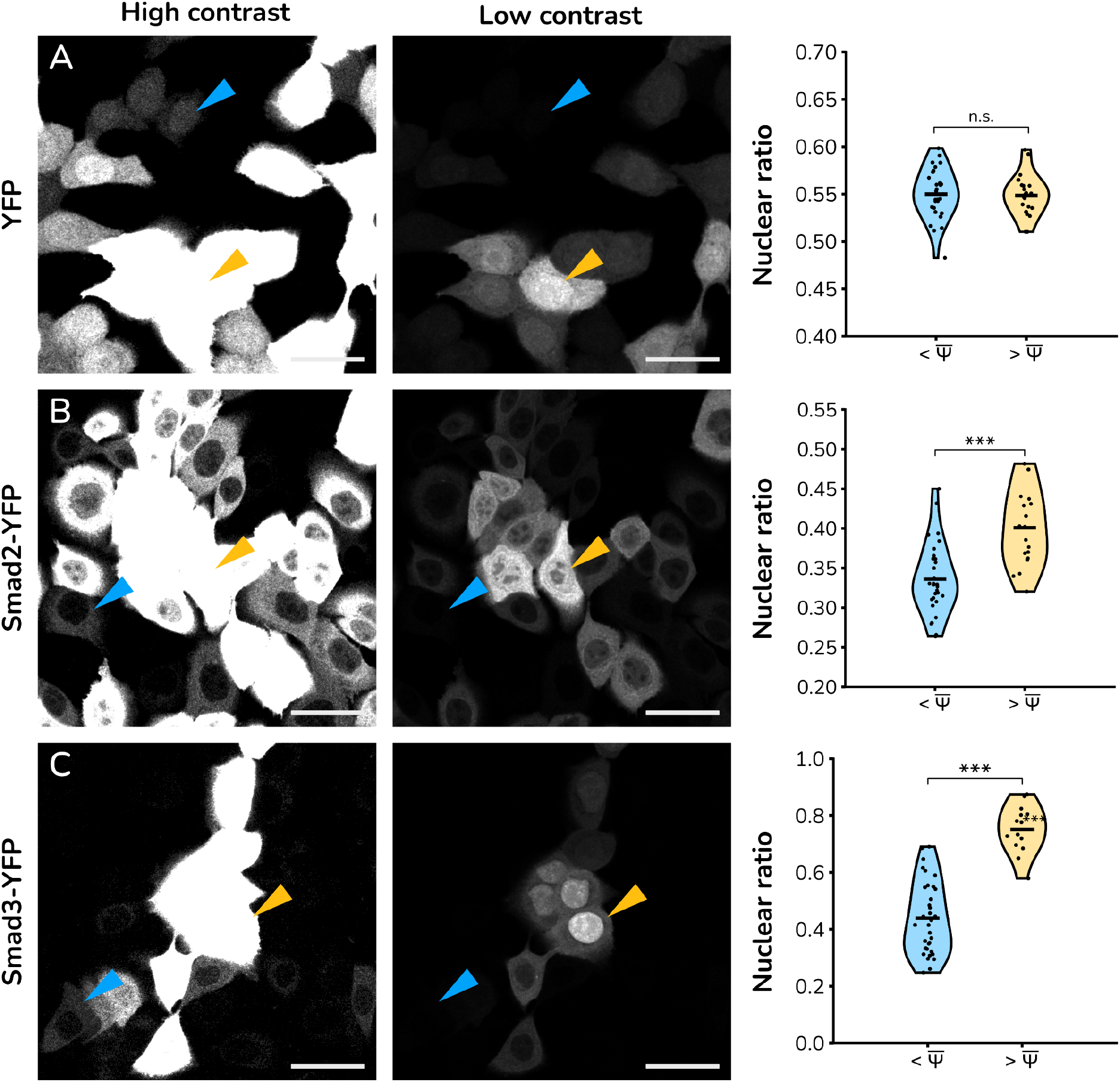
In vitro cultures of HaCaT cells overexpressing R-Smads show correlation between expression levels and nuclear ratio. Representative image of HaCaT cells stably expressing (A) YFP, (B) Smad2-YFP and (C) Smad3-YFP. In each row, the same image is represented twice for different contrast settings: increased (left panels) and decreased (central panels) levels to visualize cells expressing low and high levels of the fluorescent protein, respectively. Blue and yellow arrows point to representative cells in each image expressing low and high levels of YFP, respectively, in both images (scale bars: 50 µm). Right panels correspond to violin plots of the nuclear ratio of fluorescence, clustered by average expression levels (below and above the average fluorescence in each experiment, denoted by the value of Ψ with overscore). P-value (t-student two-tailed) is calculated to estimate statistical significance. The same data for U2OS, 293-T and 786-O are shown in Supplementary figures 1-3.

To investigate the robustness or specificity of this feature, the same experiment is carried out in cells from a highly different background. Supplementary Figures 1-3 show U2OS (human osteosarcoma), 293T (embryonic kidney), and 786-0 (renal cancer) cells infected with our lentiviral vectors. Interestingly, visual comparison suggests a higher nuclear ratio of Smad3 compared to Smad2 again, as well as the same correlation between nuclear ratio and concentration levels (right panels) of the R-Smads (absent again when overexpressing YFP alone).

In conclusion, the exogenous R-Smads show the same dependence of the nuclear ratio on the expression as the endogenous protein in tissues. The fact that the correlation between nuclear ratio and expression level occurs *in vitro* suggests that the feature arises downstream of the receptor (all cells in the *in vitro* culture are under the same upstream activation). In addition, the fact that it can be observed in cell lines with very different backgrounds and origins and basal activation levels of the TGF-β signaling pathway suggests that the process is highly general and robust.

### A mathematical model of TGF-β signaling predicts a nonlinear dependence of the nuclear ratio on the amount of protein expressed

Next, to investigate potential explanations for the correlation between expression levels and activation levels of the TGF-β pathway observed in both *in vivo* and *in vitro*, we develop a model of the signaling cascade using a mathematical formulation based on the Mass Action Law. Models of complex biological processes, such as signal regulatory networks, can be designed to incorporate all known interactions, with the potential to produce a detailed numerical picture of the system. Unfortunately, as the number of equations and free parameters increases, the complexity of these realistic models also increases, complicating their calibration and interpretation. An alternative approach is to condense biochemical steps as a single interaction, in the form of simplified conceptual models. These models are not designed to reproduce all aspects and details of the system, but to capture the underlying causes of the most relevant processes. The reduced number of equations and parameters makes them easy to interpret and calibrate with experimental data.

Following a conceptual modeling approach, we propose a generic model that focuses on the nucleocytoplasmic shuttling process, where a single protein type can enter and exit the nucleus in monomer or dimer configuration. A scheme of the numerical model is shown in Figure 4A. The specific interactions incorporated into the model are shown below:

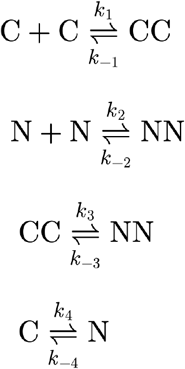

Species C and CC correspond to the concentration of protein in monomer or dimer conformation in the cytoplasm. Species N and NN correspond to the concentration of the same protein type in monomer or dimer conformation in the nucleus. First and second reactions correspond to dimerization of the protein while in the cytoplasm or in the nucleus, with binding and dissociation rates k_1_,k_-1_, and k_2_,k_-2_, correspondingly. Third and fourth reactions account for the nucleocytoplasmic shuttling as dimers and monomers, with import and export rates given by k_3_,k_-3_ and k_4_,k_-4_ correspondingly.

Mass Action Law allows us to derive the corresponding differential equations of the previous dynamical system as (detailed step-by-step derivation included as supplementary data):

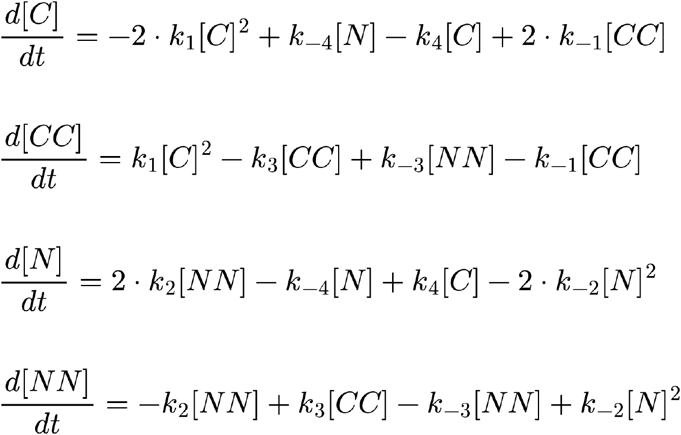

Focusing on the TGF-β pathway, it is well established that the nuclear import of monomeric Smad2 and Smad3 is negligible compared to their nuclear import rate as dimers [13, 34–37]. Similarly, the rate of nuclear export of R-Smad dimers is negligible compared to the export of monomers [13, 34–35, 38–41]. These features can be used to further simplify the model by setting k_-3_=k_4_=0. On the other hand, the association of R-Smads as dimers when in the nucleus can be considered negligible, since the kinases are at the cytoplasmic membrane, corresponding to k_2_=0 [37, 42–46]. In addition, since phosphorylated R-Smads have a very high affinity to form complexes [47], we can simplify both consecutive processes as one. This way, k_1_ can be identified as the upstream rate of activation of the pathway. Similarly, dephosphorylation of the R-Smads has been shown to trigger the disassembly of the dimer. For simplicity, the model assumes both processes as simultaneous, modulated by a single rate constant k_-2_. In addition, our previous studies show that Smad2 and Smad3 overexpression can induce 20x increase in activation levels of their transcriptional targets, evidencing that there is sufficient Smad4 to form the transcriptionally active complexes [17]. Based on this, we assume that Smad4 is in excess and does not limit the reaction kinetics. Finally, recent studies have shown that the dephosphorylation and dissociation of the active R-complexes is mediated by the CTDNEP1-NEP1R1 phosphatase complex, associated with the inner nuclear membrane protein MAN1 [48]. This confirms previous studies [37, 40, 49–50] that report that dephosphorylation and complex disassembly occurs inside the nucleus, so we can safely simplify our model by fixing the value of k_-1_=0.

After applying these simplifications, the Mass Conservation Law for the total amount of protein Ψ can be simply written as:

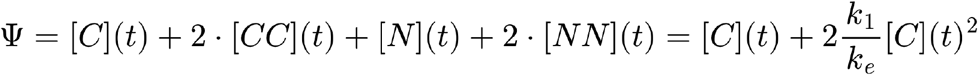

where a main relation appears to be the ratio between the activation constant k_1_ and an effective rate constant k_e_ defined as the harmonic sum of the three processes taking place sequentially (import, dissociation and export):

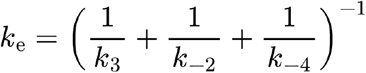

More importantly, the resulting system can be solved analytically in its steady state, providing the following equation for the nuclear concentration of R-Smad (full derivation is presented as Supplementary materials):

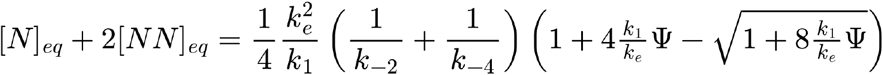

This equation predicts that the total amount of R-Smad in the nucleus at steady state depends nonlinearly on the total amount of R-Smad Ψ present in the cell.

**Figure 4.**
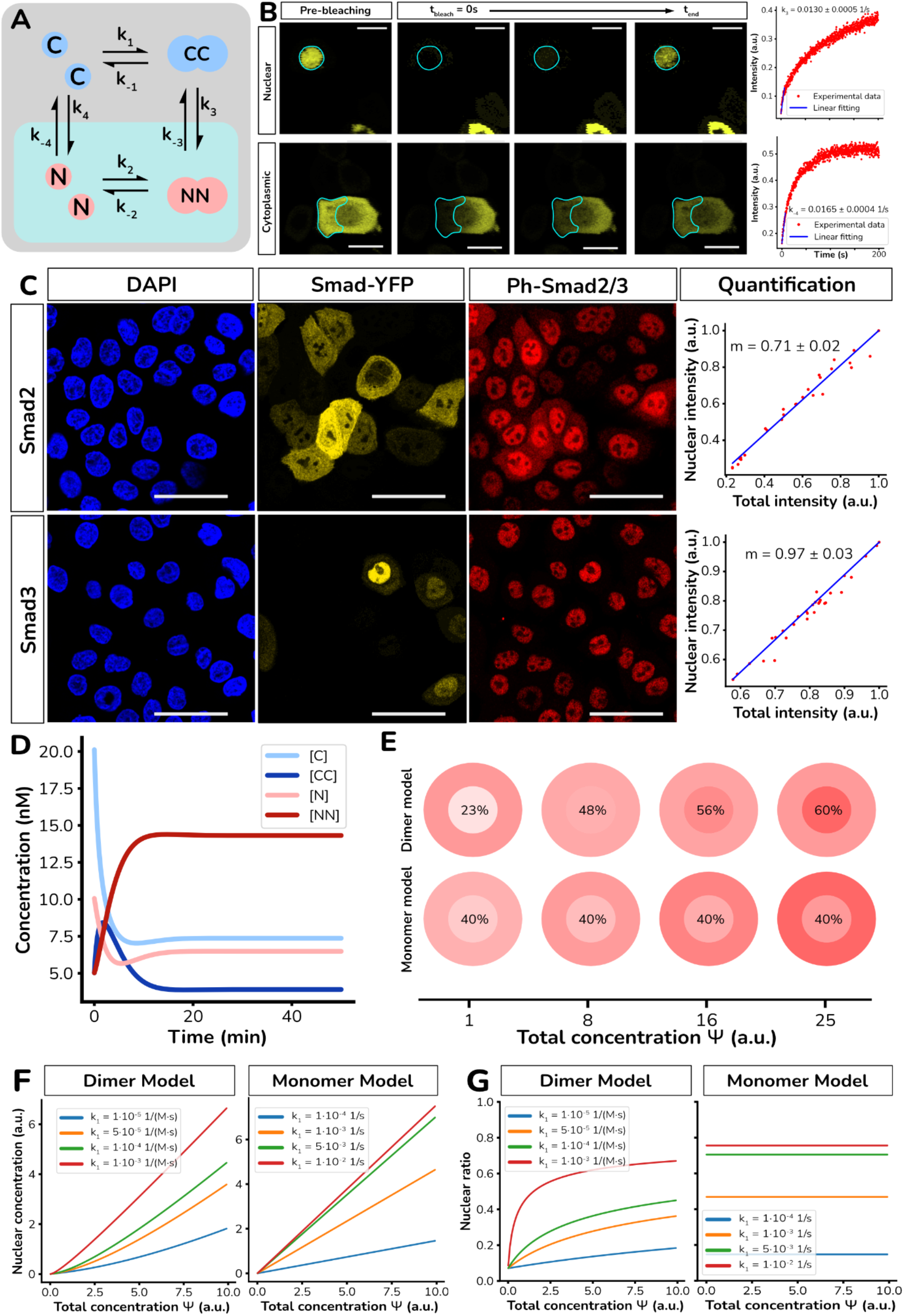
A numerical model of translocating dimers predicts correlation of nuclear ratio with expression levels. (A) Scheme of the simplified numerical model for differential nucleocytoplasmic shuttling motif. (B, upper row) Snapshots of a HaCaT cell expressing Smad3-YFP after nuclear photobleaching (scale bars: 20 µm). Contrast adjusted for visualization purposes. Quantification of temporal evolution of recovery of nuclear intensity after nuclear photobleaching to calibrate dimer import rate k_3_ is shown in the right panel. (B, lower row) Snapshots of a HaCaT cell expressing Smad2-YFP after cytoplasmic photobleaching (scale bars: 20 µm). The contrast has been adjusted for visualization purposes. Dynamics of cytoplasmic intensity after cytoplasmic photobleaching to calibrate the monomer export rate k_-4_ are shown in the right panel. (D-E) Snapshot of cells overexpressing (C) Smad2-YFP and Smad3-YFP immunostained against phospho-Smad2/3 (scale bars: 50 µm). The right panels plot the nuclear dimer ratio versus total dimer, and the linear fit (blue line) predicted by the model. (D) Numerical integration of model equations for the parameter values measured for HaCaT cells. (F) Prediction of total nuclear R-Smad levels versus total protein Ψ for different values of k_1_ in the dimer translocation and monomer translocation versions of the numerical model. (G) Prediction of nuclear ratio of R-Smad versus total protein Ψ for different values of k_1_ in the dimer translocation and monomer translocation versions of the numerical model. (E) Illustration of nuclear and cytoplasmic levels in a schematic cell predicted by the dimer translocation and monomer translocation versions of the model. Numbers indicate the value of the nuclear percentage.

Another major advantage of this simplified version of the model is that all parameters involved in k_e_ can be calibrated experimentally. To estimate the rates of import k_3_ for Smad2-YFP and Smad3-YFP, we perform Fluorescent Recovery After Photobleaching (FRAP) experiments of the nucleus of cells and estimate the value from the dynamics of nuclear fluorescence recovery. Similarly, the rates of export k_-4_ for Smad2-YFP and Smad3-YFP can be estimated by performing FRAP experiments in the cytoplasm and measuring the dynamics of cytoplasmic fluorescence recovery. Examples of these types of experiments in HaCaT cells are presented in Figure 4B as two sequences of snapshots showing nuclear FRAP and cytoplasmic FRAP. Plots of the representative dynamics of the recovery of both processes are shown in the corresponding right panels. The value of k_3_ (dimer import rate) and k_-4_ (monomer export rate) estimated by linear fitting of the initial dynamics average over independent FRAP replicas is presented in Table 1.

**Table 1:**
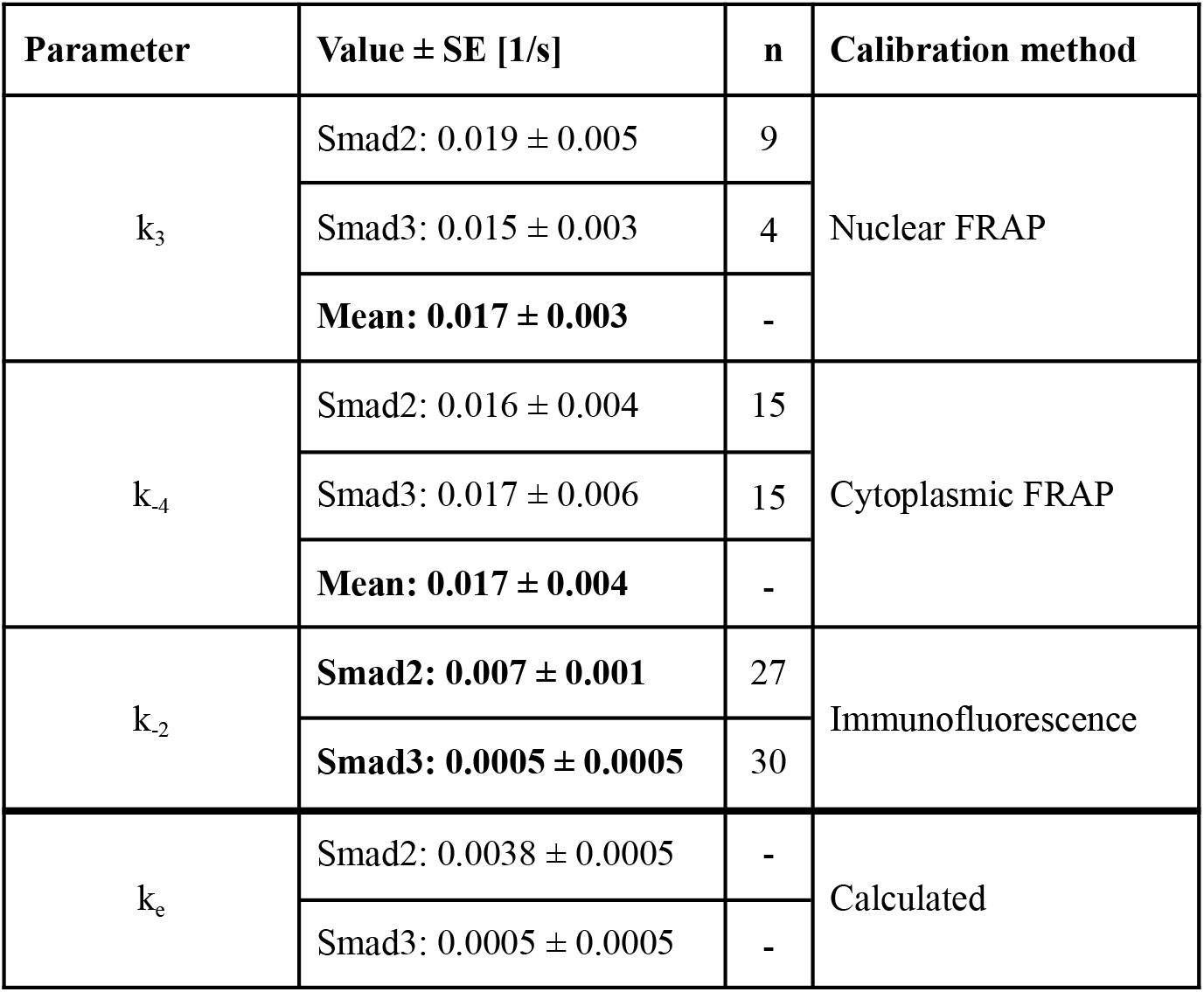
Numerical values of rate constants experimentally estimated and used for the calibration of the ODE model (in bold).

Finally, another interesting model prediction is that the nuclear concentration of dimers at equilibrium depends linearly on the total concentration of dimers in the cells (see Supplementary Material for step-by-step derivation):

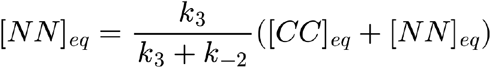

To test this prediction experimentally, we take advantage of immunostaining against the phosphorylated form of Smad2/3 (phosphorylated R-Smads have a very high affinity to form dimers). To do that, HaCaT cells overexpressing Smad2-YFP and Smad3-YFP (to induce variability in the levels of R-Smad) are stained against phospho-Smad2/3 (representative snapshots shown as Figure 4C). The right panels show a quantification of the total intensity of staining versus the nuclear intensity. As predicted by the model, the nuclear concentration of phospho-Smad2/3 increases linearly with the total concentration of phospho-Smad2/3.

Importantly, since the value of k_3_ has been measured (using nuclear FRAP), the slope of the linear regression can be used to estimate the value of k_2_. The average k_2_ value obtained after quantification of cells for each condition is shown in Table 1. This allows us to obtain the value of the effective rate constant for both Smad2 and Smad3 (Table 1). Numerical integration of the calibrated model for a constant upstream activation (represented by k_1_) is shown in Figure 4D.

For these same parameters, we plot in Figure 4F the predicted value of the total amount of R-Smad in the nucleus by the model, for different values of the total amount of protein and the activation constant k_1_. For the parameter values corresponding to HaCaT cells, the amount of R-Smad in the nucleus increases nonlinearly with the expression levels of R-Smad. For comparison, we have developed a similar model where proteins do not dimerize after activation (see Supplementary material). The solution at steady state of the nuclear concentration for this model is represented as well in Figure 4F for comparison, showing a linear dependence with the expression level, as expected.

Next, since the output of our experiments is the nuclear ratio, and since it is a widely accepted readout of TGF-β pathway activity, we plot the nuclear ratio predicted by the dimer and monomer model (Figure 4G). As expected, the monomer translocation model shows perfectly horizontal lines that change with the value of k_1_, but are independent of the amount of protein being expressed. On the contrary, the dimer model predicts that the nuclear ratio is dependent on the value of Ψ, with lower values at lower expression levels, and higher values when expression levels are high. To visually illustrate this, we represent a virtual cell where the amount of protein in the nucleus and cytoplasm is coded by color intensity. Fixing all parameter values and only changing the level of protein being present, the prediction by the dimer translocation model shows a clear visual shift in the location of the protein, with cells expressing low levels as more cytoplasmic, and cells expressing high levels as more nuclear (Figure 4E). For comparison, the same illustration is generated for the monomer model (Figure 4E), which shows a constant nuclear ratio and therefore no visual translocation of the protein.

In conclusion, a conceptual model of the differential nucleocytoplasmic translocation of monomers versus dimers calibrated for TGF-β signaling predicts a nonlinear dependence of the nuclear ratio on the amount of protein expressed. Based on this prediction, cells modulate the balance between nuclear and cytoplasmic concentration of the R-Smads depending on expression levels, just by physical processes and not requiring biochemical changes. The model correctly outputs a linear correlation between nuclear and total dimers, providing a direct validation of our approach as well as a calibration of all rates, with upstream pathway activation as the only free parameter.

### The model predicts the correlation between nuclear ratio and protein levels as a consequence of the interplay of dimerization and differential translocation

Next, to test if the model can explain the correlation between expression levels and nuclear ratio observed experimentally, we plot in Figure 5A (red dots) the nuclear ratio of Smad2 and Smad3 measured for individual cells in the HaCaT cell line, versus their total expression levels. The numerical prediction using the values for k_3_, k_-4_ and k_-2_ obtained in the previous section is superimposed (green line) over the experimental data. The value of k_1_ (upstream activation rate) for each condition and each cell line is obtained by nonlinear regression of the experimental data (see Methods). Results plotted in Figure 5A show that the nuclear ratio of YFP alone (left column) fits well the constant value predicted by the monomer translocation model. On the other hand, Smad2 (central column) goes from a cytoplasmic to a more nuclear accumulation as its expression level increases. The prediction from the dimer translocation model (green line) presents a good fitting to the experimental data.

**Figure 5:**
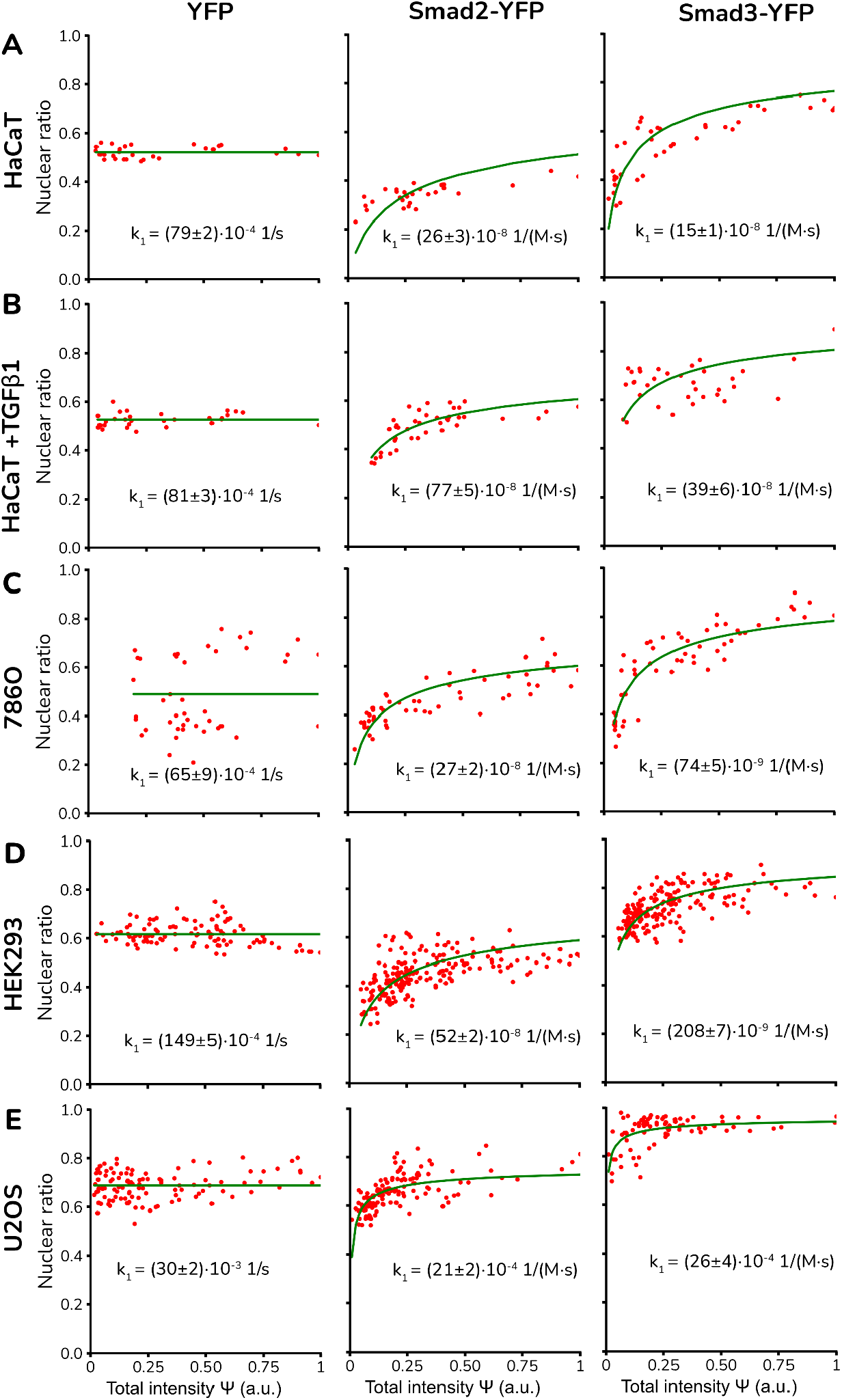
(A) Data from single-cell quantification of the HaCaT cell line overexpressing YFP (left column), Smad2-YFP (central column), and Smad3-YFP of nuclear ratio versus total expression levels. The solid line corresponds to the model solution with the activation rate k_1_ as the only fitted parameter. The monomer model has been used for YFP data, whereas the dimer model has been used for fusion proteins. (B) The same cell line is now quantified and fitted after upstream pathway activation, resulting in a higher nuclear ratio for both R-Smads and higher k_1_. (C-E) Same analysis performed for 786-O (C), HEK-293 (D), and U2OS (E) using the same calibration as the HaCaT cell line.

The nuclear ratio for Smad3 (left column) shows again a clear nonlinear dependence on its expression level. Again, the dimer model prediction using the calibrated values represents a good fit to the single-cell data. Interestingly, the value of k_1_ estimated for Smad3 is lower than for Smad2, despite sharing the same upstream activation mechanism and Smad3 appearing as more nuclear, on average. This is a consequence of the one order of magnitude difference in k_-2_ between Smad2 and Smad3.

Next, to test if our model is capable of estimating changes in the rate of upstream activation k_1_, we cultured HaCaT cells in media supplemented with TGF-β1 ligand at 50 ng/µL for one hour. Cells are then fixed, stained, quantified, and plotted in Figure 5B. As expected, the nuclear ratio of YFP is still constant and independent of the levels of TGF-β1 (left panel). On the other hand, the values of k_1_ fitted for Smad2 and Smad3 (same values for k_3_, k_-4_ and k_-2_) are increased about 2x compared to nonstimulated conditions, correlating with the stronger upstream activation due to TGF-β1 stimulation.

Next, the same experiments and quantification are repeated for the 786-O (Figure 5C), HEK293-T (Figure 5D), and U2OS (Figure 5E) cell lines. For the fitting, we assumed that the rates of import, export, and dissociation are similar to the ones measured experimentally for HaCaT cells. The quantification shows again a constant value for YFP alone (left column), and a nonconstant value for Smad2-YFP (central column) and Smad3-YFP (right column) that is well fitted by the model. Again, Smad3 is more nuclear on average than Smad2 in all cell lines tested, but the upstream activation is weaker for Smad3 in all cell lines but U2OS.

In conclusion, the increase in nuclear accumulation with the increased expression levels for both R-Smads is explained by a simplified model that is able to fit both basal activation and stimulated activation, as well as for cell lines of highly different backgrounds. This suggests that the correlation between nuclear ratio and expression levels in both R-Smads observed in all systems analyzed (*in vivo* and *in vitro*, endogenous and exogenous proteins) is a core feature of the TGF-β signaling cascade, induced by the interplay between dimerization and differential translocation of the downstream effectors of the pathway.

### Protein localization is modulated by protein expression levels in other pathways with differential dimer translocation motif

Our simplified mathematical model illustrates how a difference in the rates of nucleocytoplasmic translocation between dimers versus monomers induces coupling between protein expression and protein localization. In other words, the model suggests that any signaling cascade that contains this motif is in principle susceptible to reproducing the same feature. In fact, this nucleocytoplasmic shuttling motif is not unique to the TGF-β pathway, but a common solution present in many signaling cascades, as explained in the introduction. For instance, nucleocytoplasmic oligomer shuttling is at the core of the well-known MAPK (Mitogen-Activated Protein Kinase) pathway, involved in processes such as cell proliferation, cell differentiation, and cell death. Downstream of the pathway, ERK1 and ERK2 (Extracellular Signal-Regulated Kinase 1 and 2), undergo dimerization after phosphorylation and nuclear translocation, similarly to Smad2 and Smad3. Unlike the R-Smads, ERK1 and ERK2 do not bind directly to the DNA, and their activity as kinases does not require dimerization, so the dimerization only seems to be involved in regulating their subcellular localization.

To investigate if the ERK1/2 expression modifies its intracellular distribution, we transfected a plasmid constitutively expressing a fusion of ERK2-eGFP and fixed cells 24 hours after. Figure 6A-C shows representative images for HaCaT, U2OS, and 786-O cells, where each image is presented for high (left panel, to visualize cells expressing low levels, blue arrow) and low (central panel, to visualize cells expressing high levels, orange arrow) contrast settings. For all cases, cells expressing low levels of ERK2-eGFP have a more cytoplasmic localization than cells expressing higher amounts of the protein. The right column shows the quantification of the nuclear ratio of individual cells versus total protein expression levels (red dots). Nonlinear fitting (green line) using the model prediction is presented as a guide to show that the single-cell data fits well with a scenario where the dimerization plus translocation induces this increased nuclear ratio.

**Figure 6:**
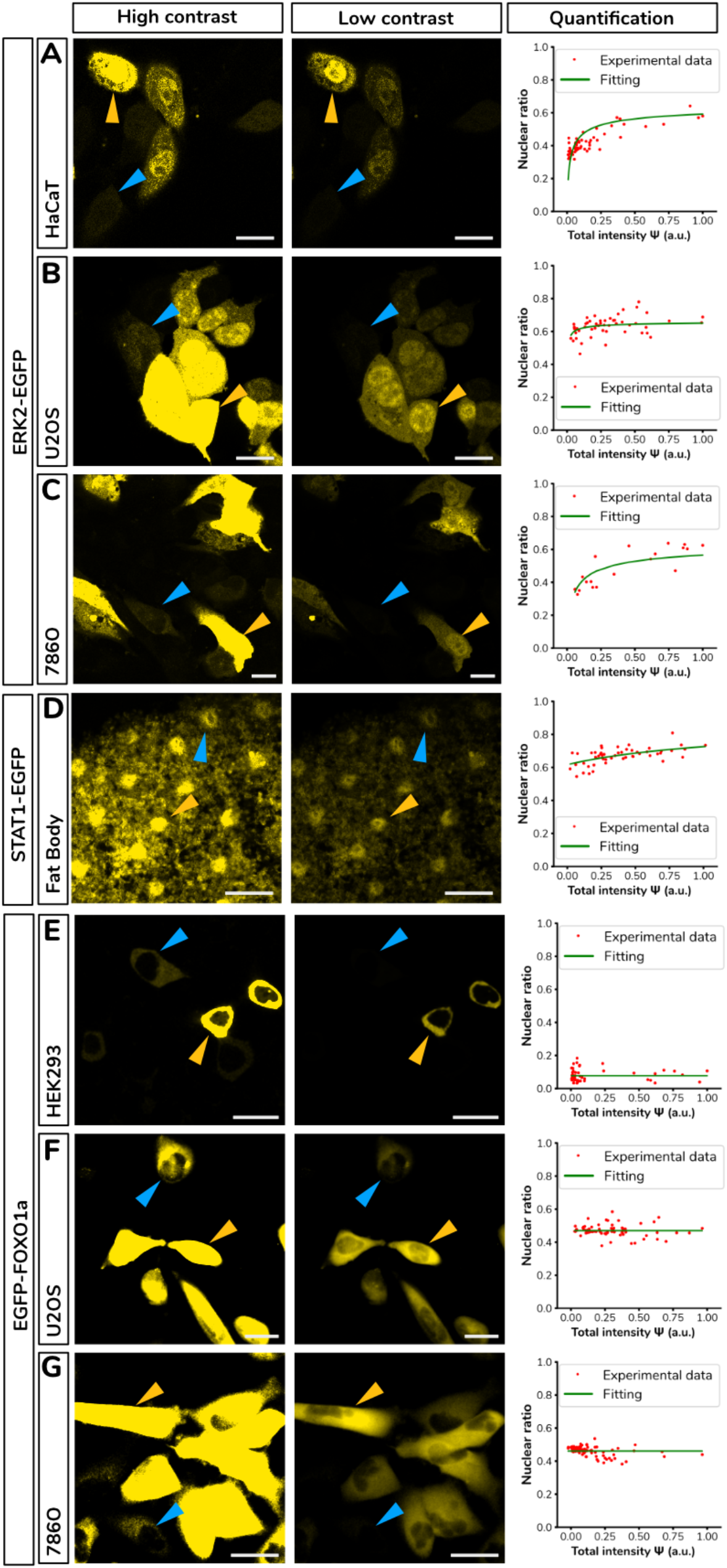
(A-C) Representative images of (A) HaCaT, (B) U2OS, and (C) 786-O cell lines overexpressing ERK2-eGFP at high (left column) and low contrast (central column). Blue and orange arrows point to cells expressing low and high levels of the protein, respectively. Plots show nuclear ratio versus total expression levels (right column) of individual cells (scale bars: 20 µm). The solid line corresponds to the dimer model fitting. (D) (Left panels) Representative image of two developing fat bodies of a Drosophila Melanogaster strain engineered to express STAT1-GFP in place of the endogenous protein. Blue and orange arrows point to cells expressing low and high levels of the protein, respectively. Plots show nuclear ratio versus total expression levels (right panel) of individual cells (scale bars: 50 µm). The solid line corresponds to the dimer model fitting. (E-G) Representative images of (E) 786-O, (F) HEK293 and (G) U2OS cell lines overexpressing FoxO1-eGFP at high (left column) and low contrast (central column). Blue and orange arrows point to cells expressing low and high levels of the protein, correspondingly. Plots show nuclear ratio versus total expression levels (right column) of individual cells (scale bars: 20 µm). The solid line corresponds to the dimer model fitting.

Another primary signaling pathway that contains the differential dimer translocation motif is the JAK/STAT signaling cascade, involved in the regulation of immune response, hematopoiesis, development, cell growth, metabolism, survival, and proliferation. Downstream of the JAK/STAT signaling is the family of Stat proteins (Signal Transducer and Activator of Transcription), which again shuttle dynamically between the nucleus and cytoplasm depending on phosphorylation-driven dimerization, sensing signals from the cell membrane and physically translating into regulating specific genes. To investigate the intracellular organization of STAT, we take advantage of a fruit fly strain engineered to express STAT1 (the only member of the STAT family in flies) fused to eGFP, introduced directly by replacing the locus of the STAT1 gene. This allows us to test the effect of the differential dimer translocation motif in a context of endogenous protein levels. In this system, we focus on the fat body tissue of developing fly embryos because its cells exhibit sufficiently high expression levels of STAT1, and their large cytoplasm facilitates the quantification of the nuclear ratio, compared to other tissues such as the wing or eye disc. Figure 6D shows representative images of this strain to visualize the boundaries of each individual cell. Visual inspection of the staining shows cells expressing different levels of STAT1-eGFP (marked with blue arrows for low levels of expression and orange arrows for high levels, correspondingly). The left panel plots the nuclear ratio versus expression levels, with the curve predicted by the model (solid line) representing a good fit to the data.

Next, we focus on the AKT signaling cascade, another central pathway in cellular biology that heavily relies on protein translocation for the regulation of its targets. FoxO1 is a key transcription factor directly downstream of AKT that translocates to the nucleus depending on its phosphorylation state. The main difference with the R-Smads, Stat, or Herk families is that Foxo proteins do not form oligomers. Again, we proceed to transfect a plasmid constitutively expressing a fusion of FoxO1-eGFP into 786-O (Figure 6E), HEK293 (Figure 6F) and U2OS (Figure 6G). Images of the cells 24 hours after transfection (again, low- and high-contrast versions of the same image are included) show that the expression levels of the fusion FoxO1-eGFP are highly variable, but most of the cells show a similar organization in terms of the nuclear ratio of the fluorescence levels. Quantification (right panels) shows that the nuclear ratio of FoxO1 is independent of its expression level (fitting corresponding with the monomer model is included, green line).

In conclusion, all tested signaling pathways with a differential dimer translocation motif (JAK/STAT, MAPK and TGF-β) show the same correlation between expression levels and nuclear ratio of their main effector proteins. This feature is not present in the AKT pathway, since the nucleocytoplasmic shuttling does not involve dimerization, as predicted by our model.

## Discussion

The activation of signal-transduction pathways is often depicted as triggered by extracellular ligands that activate receptors that activate cytoplasmic transducers that activate transcription factors that ultimately activate (or repress) gene expression. All the accepted seven major [51] signaling pathways (Wnt, TGF-β, Hh, RTK–MAPK, JAK/STAT, Notch, and nuclear receptor) incorporate a mechanism of physical separation of active versus inactive effector proteins (β-Catenin, R-Smads, GLI, ERK1/2, STAT1/2, Notch Intracellular Domain, and Hsp90) achieved via different rates of nuclear import/export. In three of them, this compartmentalization occurs as active dimers versus inactive monomers (TGF-β, RTK–MAPK and JAK/STAT). Here we show that if monomers and dimers have different nuclear import/export rates, the combination of differential translocation and dimerization acts as a network motif that couples expression levels and nuclear ratio.

Another potential explanation will involve ligands activating TGF-β expressed in the same regions as the signals that drive expression of R-Smads, and this occurring similarly in such diverse tissues as the spinal cord, in the human brain, and also in early developing mammalian embryos (well-characterized signals involved in the regulation of Smad2 are p53-MEG3 [52], and JAK/STAT3 for Smad3 [53]). Also, a potential positive feedback loop between protein levels and activation can, in principle, produce a similar effect, but it is widely accepted that there are no positive feedbacks involved in the TGF-β signaling cascade [54–58]. Finally, another potential explanation is a potential saturation of the nuclear export machinery, i.e., resulting in a reduction of the rate of export when the number of R-Smad molecules is high. To rule this out, we measured the export rate (k_-4_) in individual cells using FRAP (see Methods) and observed no dependence of this rate on the expression level (see Supplementary Figure 4). Also, the fact that the effect occurs for endogenous levels (for both TGF-β and Stat pathways), and that the effect is absent in conditions of overexpression of FoxO1, suggests again that the effect is not due to nuclear export saturation. Finally, this scenario of saturation of nuclear export is tested numerically using a variation of the linear model where k_-4_ is modulated by a repressing Hill function modulated by the amount of molecule to be exported (N in this case; details described in the supplementary information). The nuclear ratio predicted is shown in Supplementary Figure 5AB for n=1 (no cooperativity) and n=2 (cooperativity). In both cases, even for a Hill parameter as low as 1 (saturation is taking place at around 100 times less than the expression level), the export saturation model (red curves) predicts nuclear ratios very different from the curves predicted by the differential dimer translocation model (blue curve).

Focusing on the mechanism underneath the coupling, the linear model (Figure 4G, similar to the FoxO1 translocation experiments, Figure 6D-G) illustrates that dimerization is a key component of the motif. To study the effect of the other ingredient (differential rates), we generated a version of the model with similar import and export rates for dimers and monomers. In this model, the nuclear ratio (shown in Supplementary Figure 5C) is again independent of the concentration, evidencing that both ingredients (dimerization and differential translocation) are essential parts of the motif.

Previous studies (including our own [17]) identify regions of higher R-Smad nuclear ratio with regions where the upstream activation of the pathway is higher. Here we show that an increase in R-Smad expression increases its nuclear ratio independently of upstream pathway levels. A key difference between the mechanism of differential nucleocytoplasmic translocation of dimers versus monomers and other previously known network motifs is that it does not involve a loop of autoregulation (feedback or feedforward), and that it introduces a nonlinear dependence using intracellular physical compartments.

## Conclusions

Here, we present a network motif operation in signaling pathways based on differential nucleocytoplasmic translocation of monomers versus dimers, introducing a nonlinear coupling between expression levels of a protein and its intracellular translocation. This mechanism modulates activity of the TGF-β signaling cascade by affecting the location of its main effectors, the R-Smads, and explains the increase in nuclear ratio in regions of higher expression observed in all tissues and cell lines analyzed in this study. Moreover, the network motif seems to be operating also in all signaling cascades analyzed that incorporate a dimerization step linked to nucleocytoplasmic shuttling, operating as a singular type of nonlinear network motif that utilizes compartmentalization and does not require a loop of regulation.

## Acknowledgments

This work was supported by grants from the Ministerio de Ciencia e Innovación, Spain [RTI2018-096953-B-I00, PID2022-140421NB-I00 and PDC2022-133147-I00, to D.G.M.; PREP2022-000149 founded by MCIN/AEI/10.13039/501100011033 and EFS+ to DM-D, and Institutional fellowships to IFIMAC (Maria de Maeztu Unit of Excellence; CEX2023-001316-M) and CBMSO (Severo Ochoa Unit of Excellence)]. We acknowledge all the Center for Molecular Biology Severo Ochoa (CBMSO) facilities, including the Advanced Optical Microscopy Service (SMOA) and the Animal facility.

## Notes

### Competing Interest Statement

The authors have declared no competing interest.

