## Supplementary Information for "A signaling network motif couples protein expression levels and intracellular localization"

### Conceptual translocation model

This simple model captures the core dynamics of one of the most common patterns in cell biology. In many processes, monomeric proteins in the cell cytoplasm must aggregate to form complexes that then translocate into the nucleus. There, the complexes are disassembled, and the monomers come back to the cytoplasm to start the cycle again. To simplify, for this model we will consider that complexes are just made up of two monomeric proteins; that is, they form dimers.

Let C be a cytoplasmic monomeric protein which can dimerize into the dimer CC. This dimerization process has a rate $k_1$, and as it can be reversible, we will consider that they can separate back into two monomers at a rate $k_{-1}$. Both monomers C and dimers CC can translocate to the nucleus of the cell, becoming N and NN, respectively. These translocation processes are reversible too, so if CC can enter into the nucleus to become NN at a rate $k_3$, NN can exit the nucleus at a rate $k_{-3}$. The same is true for C becoming N at a rate $k_4$ (nuclear import) and N becoming C at a rate $k_{-4}$ (nuclear export). Inside the nucleus, nuclear monomers N can form dimers NN at a rate $k_2$, and dimers can split into monomers at a rate $k_{-2}$ (it is a reversible process). We can summarize all these interactions into the following chemical reactions:

$$\mathrm{C} + \mathrm{C} \underset{k_{-1}}{\overset{k_1}{\rightleftharpoons}} \mathrm{CC}$$

$$\mathrm{N} + \mathrm{N} \underset{k_{-2}}{\overset{k_2}{\rightleftharpoons}} \mathrm{NN}$$

$$\mathrm{CC} \underset{k_{-3}}{\overset{k_3}{\rightleftharpoons}} \mathrm{NN}$$

$$\mathrm{C} \underset{k_{-4}}{\overset{k_4}{\rightleftharpoons}} \mathrm{N}$$

Now, we can use the Mass Action Law to obtain a set of ordinary differential equations (ODE) which model the dynamics of this system over time. An in-depth explanation of the mathematical procedure to obtain these equations can be found in **[Cellular Compartmentalization as a Physical Regulatory Mechanism of Signaling Pathways]**. For that, let first [C], [N], [CC], and [NN] be the concentrations of monomers in the cytoplasm and the nucleus and the concentrations of dimers in the cytoplasm and the nucleus, respectively. Let A and B be two matrices, containing the stoichiometric coefficients of the reactants and products, respectively. Finally, let [\vec{X}] be a column vector containing the concentrations of the species and let K be a diagonal matrix with the reaction rates. In this particular context, these mathematical objects are explicitly defined as:

$$[\vec{X}] = \begin{pmatrix} [C] \\ [CC] \\ [N] \\ [NN] \end{pmatrix}$$

$$K = \begin{pmatrix} k1 & 0 & 0 & 0 & 0 & 0 & 0 & 0 \\ 0 & k_{-1} & 0 & 0 & 0 & 0 & 0 & 0 \\ 0 & 0 & k_2 & 0 & 0 & 0 & 0 & 0 \\ 0 & 0 & 0 & k_{-2} & 0 & 0 & 0 & 0 \\ 0 & 0 & 0 & 0 & k_3 & 0 & 0 & 0 \\ 0 & 0 & 0 & 0 & 0 & k_{-3} & 0 & 0 \\ 0 & 0 & 0 & 0 & 0 & 0 & k_4 & 0 \\ 0 & 0 & 0 & 0 & 0 & 0 & 0 & k_{-4} \end{pmatrix}$$

$$A = \begin{pmatrix} 2 & 0 & 0 & 0 \\ 0 & 1 & 0 & 0 \\ 0 & 0 & 0 & 0 \\ 0 & 0 & 2 & 0 \\ 0 & 1 & 0 & 0 \\ 0 & 0 & 0 & 1 \\ 1 & 0 & 0 & 0 \\ 0 & 0 & 1 & 0 \end{pmatrix}$$

$$B = \begin{pmatrix} 0 & 1 & 0 & 0 \\ 2 & 0 & 0 & 0 \\ 0 & 0 & 2 & 0 \\ 0 & 0 & 0 & 1 \\ 0 & 0 & 0 & 1 \\ 0 & 1 & 0 & 0 \\ 0 & 0 & 1 & 0 \\ 1 & 0 & 0 & 0 \end{pmatrix}$$

Using the definition of the Mass Action Law

$$\frac{dX}{dt} = (B - A)^t \cdot K \cdot X^A$$

we can obtain the desired ODE system:

$$\frac{d[C]}{dt} = -2k_1[C]^2 + 2k_{-1}[CC] - k_4[C] + k_{-4}[N]$$
$$\frac{d[CC]}{dt} = k_1[C]^2 - k_{-1}[CC] - k_3[CC] + k_{-3}[NN]$$
$$\frac{d[N]}{dt} = 2k_{-2}[NN] - 2k_2[N]^2 + k_4[C] - k_{-4}[N]$$
$$\frac{d[NN]}{dt} = -k_{-2}[NN] + k_2[N]^2 + k_3[CC] - k_{-3}[NN]$$

For this system, the Mass Conservation Law is:

$$[C] + 2[CC] + [N] + 2[NN] = \Psi \quad (1)$$

where $\psi$ is a constant that represents the total concentration of monomer proteins in the cell. This value remaining constant implies that no protein is being produced nor degraded.

### TGF-β translocation model

Once we have obtained the general equations for the translocation dynamics, we can apply them to the specific case of the Smad2-Smad3 TGF-β pathway. To do so, let's make some simplifications and assumptions. Firstly, we will consider that Smad4 is not a limiting reagent, so we can just focus on the modeling of the Smad2 and Smad3 proteins. Secondly, as these R-Smads are quite similar, we will consider them as the same molecule. With these assumptions, C and N represent Smad (2 and/or 3) monomers in the cytoplasm and the nucleus, respectively; and CC and NN represent Smad (2-2, 2-3 or 3-3) complexes (not considering Smad4) in the cytoplasm and nucleus, respectively.

On top of that, we will assume that monomeric Smads do not dimerize in the nucleus and dimers do not separate in the cytoplasm, thus yielding $k_{-1} = k_2 = 0$. Moreover, if we consider that monomers cannot enter the nucleus and dimers can not exit this compartment, then $k_{-3} = k_4 = 0$. With all these simplifications, our TGF-β model ODE system remains as:

$$\frac{d[C]}{dt} = -2k_1[C]^2 + k_{-4}[N]$$

$$\frac{d[CC]}{dt} = k_1[C]^2 - k_3[CC]$$
$$\frac{d[N]}{dt} = 2k_{-2}[NN] - k_{-4}[N]$$
$$\frac{d[NN]}{dt} = -k_{-2}[NN] + k_3[CC]$$

Note that all rates are measured in 1/s except for $k_1$, which is measured in 1/(M·s).

<u>Values in the steady state</u>

With this ODE system, we can get the concentrations in the steady state for the different species. First, we set each derivative to zero (definition of steady state), so that we get a system of linear equations. Then, we can express each concentration in terms of one of them (in this case, we choose [C] arbitrarily):

$$0 = -2k_1[C]^2 + k_{-4}[N] \qquad (2)$$
$$0 = k_1[C]^2 - k_3[CC] \qquad (3)$$
$$0 = 2k_{-2}[NN] - k_{-4}[N] \qquad (4)$$
$$0 = -k_{-2}[NN] + k_3[CC] \qquad (5)$$

From Equation 2 and Equation 3, we can get to:

$$[N] = 2\frac{k_1}{k_{-4}}[C]^2 \qquad (6)$$
$$[CC] = \frac{k_1}{k_3}[C]^2 \qquad (7)$$

Using Equation 5 and Equation 3, we have:

$$k_{-2}[NN] = k_3[CC] = k_1[C]^2 \rightarrow [NN] = \frac{k_1}{k_{-2}}[C]^2 \qquad (8)$$

As we now have all the species in terms of one of them, we can use the Mass Conservation Law stated in Equation 1 to reach an expression of [C] as a function of $\psi$:

$$\Psi = [C] + 2[CC] + [N] + 2[NN] = [C] + 2\frac{k_1}{k_3}[C]^2 + 2\frac{k_1}{k_{-4}}[C]^2 + 2\frac{k_1}{k_{-2}}[C]^2$$

$$\Psi = [C] + 2k_1[C]^2 \left( \frac{1}{k_{-2}} + \frac{1}{k_3} + \frac{1}{k_{-4}} \right)$$

$$[C] = \frac{\sqrt{1 + 8\frac{k_1}{k_e}\Psi} - 1}{4\frac{k_1}{k_e}} \quad (9)$$

where we define the effective rate $k_e$ as

$$\frac{1}{k_e} = \frac{1}{k_{-2}} + \frac{1}{k_3} + \frac{1}{k_{-4}}$$

which encapsulates the rate constants that can be experimentally calibrated, as we will explain later in this document. For now, using Equation 9 and Equation 6, Equation 7 and Equation 8, we can get the value of the concentration of all the species in the steady state:

$$[CC] = \frac{1}{8}\frac{k_e^2}{k_1 k_3} \cdot \left(1 + 4\frac{k_1}{k_e}\Psi - \sqrt{1 + 8\frac{k_1}{k_e}\Psi}\right)$$

$$[N] = \frac{1}{4}\frac{k_e^2}{k_1 k_{-4}} \cdot \left(1 + 4\frac{k_1}{k_e}\Psi - \sqrt{1 + 8\frac{k_1}{k_e}\Psi}\right)$$

$$[NN] = \frac{1}{8}\frac{k_e^2}{k_1 k_{-2}} \cdot \left(1 + 4\frac{k_1}{k_e}\Psi - \sqrt{1 + 8\frac{k_1}{k_e}\Psi}\right)$$

<u>Smad nuclear ratio in the steady state</u>

With all the mathematical expressions that we have derived above, we are now able to calculate which ratio of the total Smad concentration $\psi$ within a cell is expected to be retained in the nucleus depending on that total Smad concentration $\psi$. Mathematically, this ratio is defined as:

$$\frac{[N] + 2[NN]}{\Psi}$$

Using Equation 6, Equation 8 and Equation 9:

$$\frac{[N] + 2[NN]}{\Psi} = \frac{1}{4}\frac{k_e^2}{k_1\Psi}\left(\frac{1}{k_{-2}} + \frac{1}{k_{-4}}\right)\left(1 + 4\frac{k_1}{k_e}\Psi - \sqrt{1 + 8\frac{k_1}{k_e}\Psi}\right)$$

Smad dimer nuclear ratio in the steady state

Another quantity of interest that our model can predict is the ratio of Smad dimers that reside in the nucleus. Mathematically, it is expressed as

$$\frac{[NN]}{[CC]+[NN]}$$

where [CC]+[NN] represents the total amount of dimers in the cell. Thanks to Equation 7 and Equation 8, we get to:

$$\frac{[NN]}{[CC]+[NN]} = \frac{k_3}{k_3+k_{-2}}$$

Note that this ratio does not depend on the total concentration of proteins $\psi$.

### Linear translocation model

Now, let's derive the equations for another common situation in which monomers do not need to dimerize to translocate into the nucleus, but they do need to activate somehow. Biologically, this could be a phosphorylation process or simply a conformational change. The core of the system remains unaltered, but we will change CC and NN by C* and N*, indicating that C and N have been activated in any manner. Now, $k_1$ and $k_{-1}$ become the activation and deactivation rates for cytoplasmic monomers, and $k_2$ and $k_{-2}$ for nuclear monomers, respectively. Translocation rates behave similarly: $k_3$ and $k_4$ refer to the nuclear import rates for C* and C, respectively; and $k_{-3}$ and $k_{-4}$ refer to the nuclear export rates for N* and N, respectively. With all these considerations, the set of chemical reactions that describes this system is:

$$C \underset{k_{-1}}{\overset{k_1}{\rightleftharpoons}} C^*$$

$$N \underset{k_{-2}}{\overset{k_2}{\rightleftharpoons}} N^*$$

$$C^* \underset{k_{-3}}{\overset{k_3}{\rightleftharpoons}} N^*$$

$$C \underset{k_{-4}}{\overset{k_4}{\rightleftharpoons}} N$$

Analogously to what we did above, using the Mass Action Law, we can reach the ODE system that describes the dynamics of this situation. For it to be comparable with the TGF-β model, we will also consider that $k_{-1} = k_2 = k_{-3} = k_4 = 0$:

$$\frac{d[C]}{dt} = -k_1 C + k_{-4} N$$
$$\frac{d[C^*]}{dt} = k_1 C - k_3 C^*$$
$$\frac{d[N]}{dt} = k_{-2} N^* - k_{-4} N$$
$$\frac{d[N^*]}{dt} = -k_{-2} N^* + k_3 C^*$$

Note that, in this case, the Mass Conservation Law is written as:

$$\Psi = [C] + [C^*] + [N] + [N^*]$$

and in this case, all constant rates are expressed in 1/s.

Again, if we consider the steady state, similarly to what we did above, we obtain the expressions of the species as a function of $\psi$:

$$[C] = \frac{\Psi k_e}{k_1 + k_e}$$
$$[C^*] = \frac{k_1}{k_3}[C] = \frac{\Psi k_1 k_e}{k_3(k_1 + k_e)}$$
$$[N] = \frac{k_1}{k_{-4}}[C] = \frac{\Psi k_1 k_e}{k_{-4}(k_1 + k_e)}$$
$$[N^*] = \frac{k_1}{k_{-2}}[C] = \frac{\Psi k_1 k_e}{k_{-2}(k_1 + k_e)}$$

Furthermore, we can also calculate the ratio of proteins in the nucleus at that steady state:

$$\frac{[N] + [N^*]}{\Psi} = \frac{k_1 k_e}{k_1 + k_e} \cdot \left( \frac{1}{k_{-4}} + \frac{1}{k_{-2}} \right)$$

and the nuclear ratio of active monomers in the steady state:

$$\frac{[N^*]}{[N^*] + [C^*]} = \frac{k_3}{k_3 + k_{-2}}$$

Up to this point, it is worth noticing that the ratio of proteins in the nucleus in the steady state does not depend on $\psi$, and that the nuclear ratio of active monomers in the steady state is the same as the one predicted by the TGF-β model (and independent of $\psi$, too).

Negative feedback on $k_{-4}$ depending on [N]

We developed a modified version of the linear model including a negative feedback depending on [N], modifying the value of $k_{-4}$. To do so, we start from the linear model ODE system and multiply $k_{-4}$ in every equation by a negative Hill function defined as

$$h_{k,r}([N]) = \frac{k^r}{k^r + [N]^r}$$

where k is a constant and r is the Hill coefficient. This modification implies that the higher the concentration of [N], the less it is exported to the cytoplasm, leading to a nuclear accumulation of the protein. Essentially, this introduces a saturation dynamic into the model. With this new ODE system, we can obtain the nuclear ratio of protein as a function of $\psi$ numerically, as shown in Supplementary Figure 5AB.

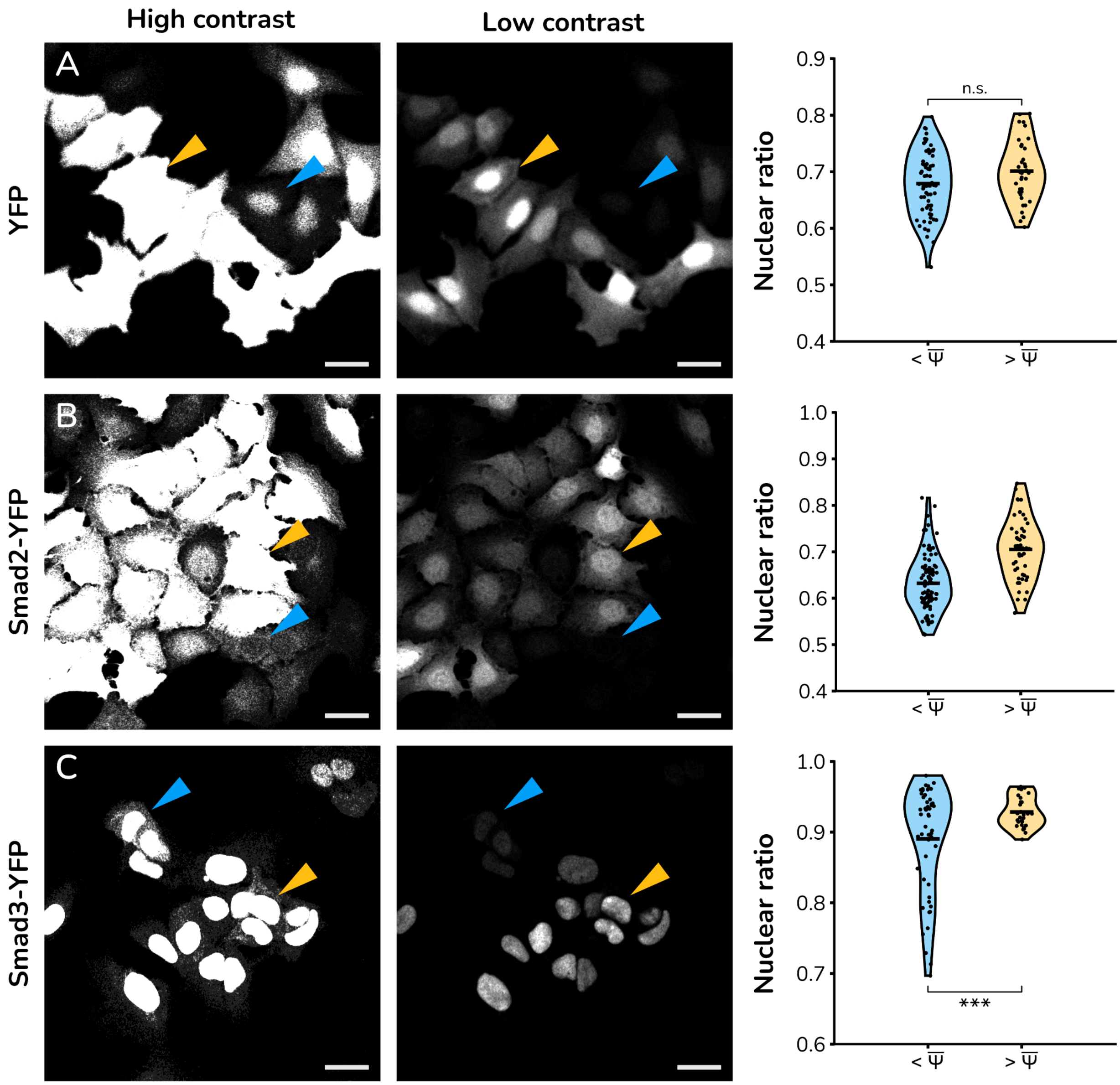


***Supplementary Figure 1.*** *In vitro cultures of U2OS cells overexpressing R-Smads show correlation between expression levels and nuclear ratio. Representative image of HaCaT cells stably expressing (A) YFP, (B) Smad2-YFP and (C) Smad3-YFP. In each row, the same image is represented twice for different contrast settings: increased (left panels) and decreased (central panels) levels to visualize cells expressing low and high levels of the fluorescent protein, respectively. Blue and yellow arrows point to representative cells in each image expressing low and high levels of YFP, respectively, in both images (scale bars: 50 µm). Right panels correspond to violin plots of the nuclear ratio of fluorescence, clustered by average expression levels (below and above the average fluorescence in each experiment). P-value (t-test two-tailed) is calculated to estimate statistical significance.*

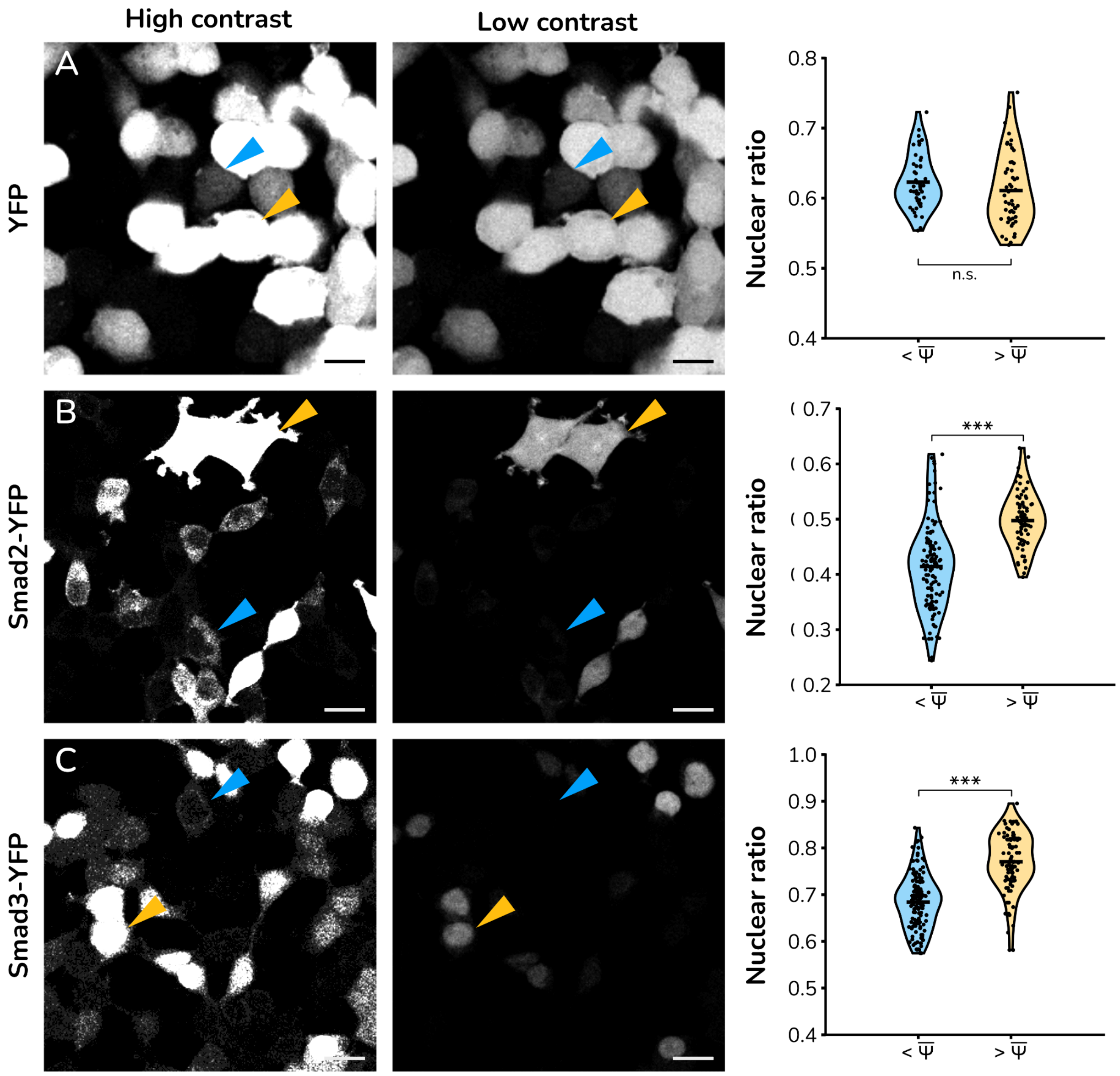


***Supplementary Figure 2.*** *In vitro cultures of HEK293 cells overexpressing R-Smads show correlation between expression levels and nuclear ratio. Representative image of HaCaT cells stably expressing (A) YFP, (B) Smad2-YFP and (C) Smad3-YFP. In each row, the same image is represented twice for different contrast settings: increased (left panels) and decreased (central panels) levels to visualize cells expressing low and high levels of the fluorescent protein, respectively. Blue and yellow arrows point to representative cells in each image expressing low and high levels of YFP, respectively, in both images (scale bars: 50 µm). Right panels correspond to violin plots of the nuclear ratio of fluorescence, clustered by average expression levels (below and above the average fluorescence in each experiment). P-value (t-test two-tailed) is calculated to estimate statistical significance.*

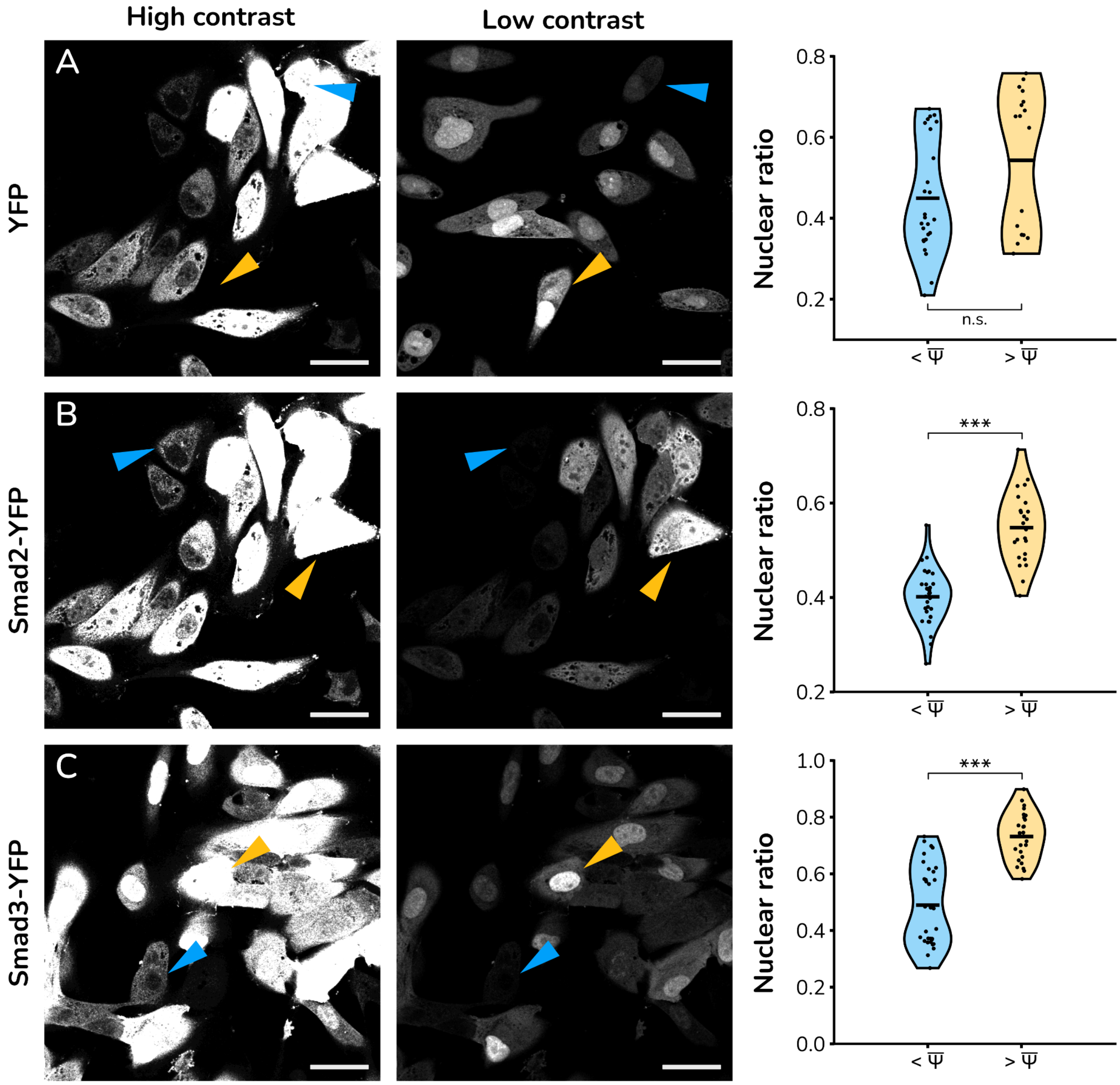


***Supplementary Figure 3.*** *In vitro cultures of 786-O cells overexpressing R-Smads show correlation between expression levels and nuclear ratio. Representative image of HaCaT cells stably expressing (A) YFP, (B) Smad2-YFP and (C) Smad3-YFP. In each row, the same image is represented twice for different contrast settings: increased (left panels) and decreased (central panels) levels to visualize cells expressing low and high levels of the fluorescent protein, respectively. Blue and yellow arrows point to representative cells in each image expressing low and high levels of YFP, respectively, in both images (scale bars: 50 µm). Right panels correspond to violin plots of the nuclear ratio of fluorescence, clustered by average expression levels (below and above the average fluorescence in each experiment). P-value (t-test two-tailed) is calculated to estimate statistical significance.*

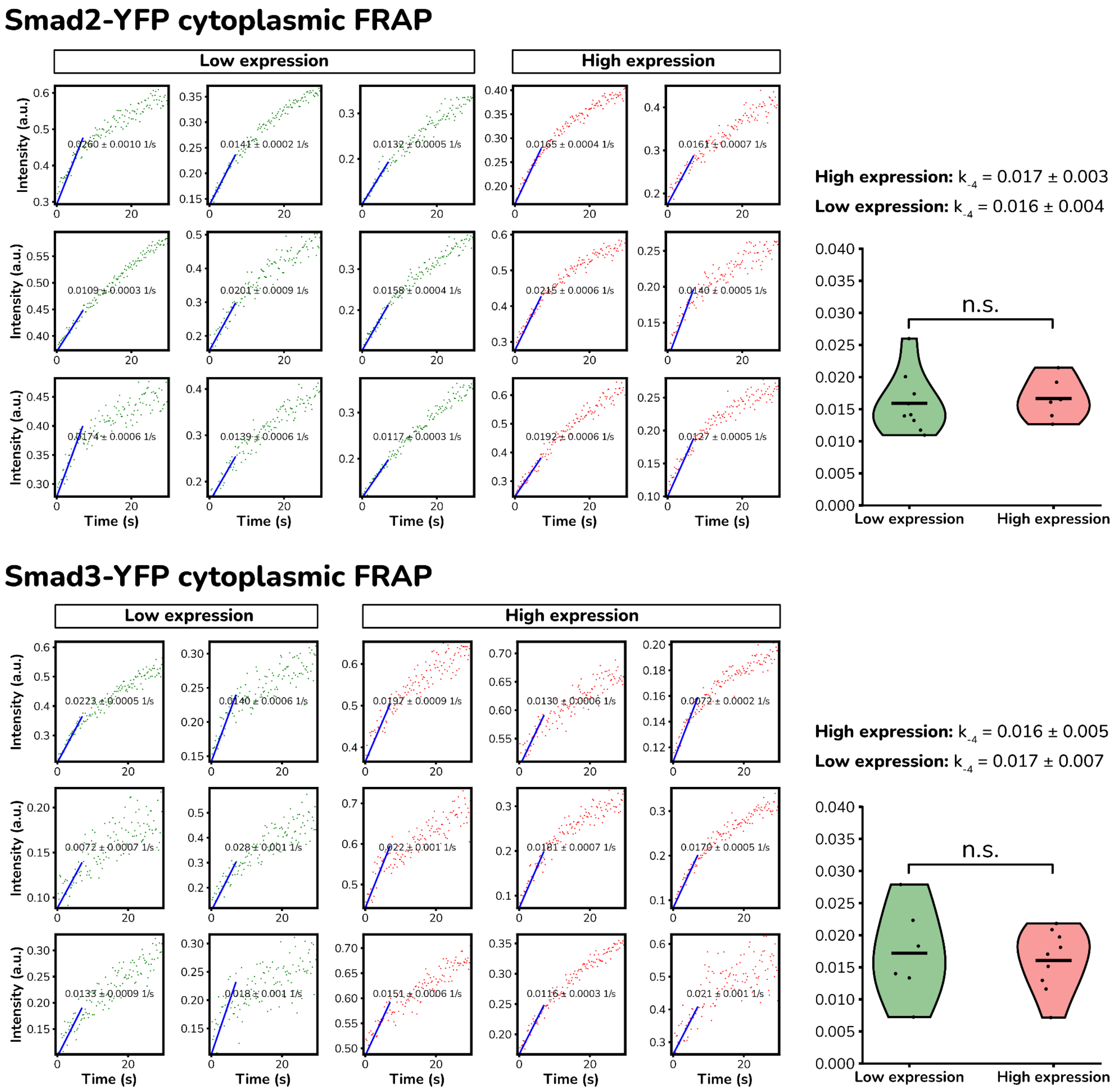


***Supplementary Figure 4.*** *Dynamic Data of Single cells after FRAP clustered by cells expressing below-average (green) and above-average (red) levels of YFP-Smad2 (above) and YFP-Smad3. Plots in the right column show the quantification of the value of* $k_{-4}$ *with nonsignificant statistical differences between the two datasets,*

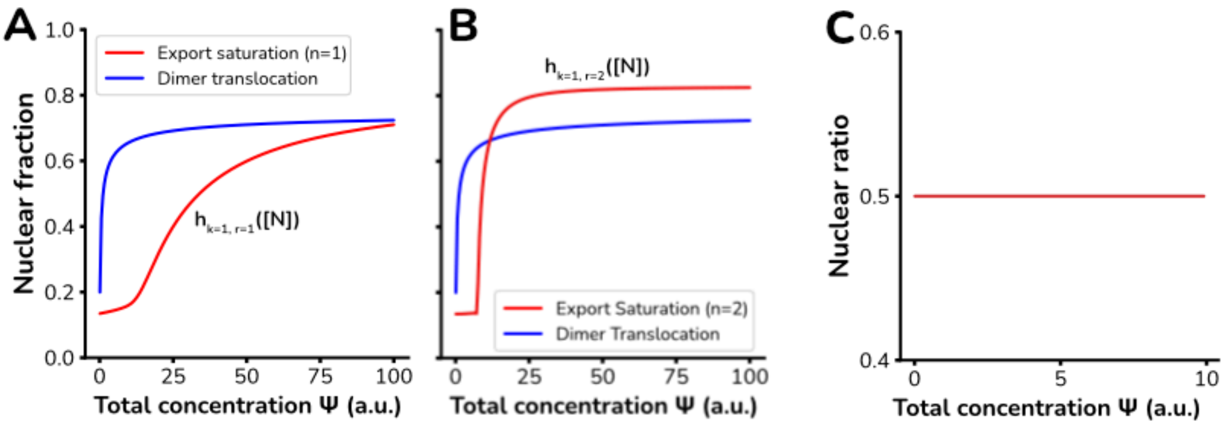


***Supplementary Figure 5.*** ***(A)*** *Nuclear ratio predicted by the export saturation model (red line) compared to the dimer model (blue line) for non-cooperativity Hill function (left panel) and cooperativity (n=2, right panel).* ***(B)*** *Nuclear ratio predicted by the dimer model with non-differential nucleocytoplasmic translocation between monomers and dimers.*
